# Pro-inflammatory polarization and colorectal cancer modulate alternative and intronic polyadenylation in primary human macrophages

**DOI:** 10.1101/2023.02.24.529734

**Authors:** Joana Wilton, Filipa Lopes de Mendonça, Isabel Pereira-Castro, Michael Tellier, Takayuki Nojima, Angela M Costa, Jaime Freitas, Shona Murphy, Maria Jose Oliveira, Nicholas J Proudfoot, Alexandra Moreira

**Affiliations:** Graduate Program in Areas of Basic and Applied Biology (GABBA) PhD Program, ICBAS-Instituto de Ciências Biomédicas Abel Salazar, Universidade do Porto, Portugal; Gene Regulation - Instituto de Investigação e Inovação em Saúde, Universidade do Porto, Portugal; IBMC-Instituto de Biologia Molecular e Celular, Portugal; Sir William Dunn School of Pathology, University of Oxford, United Kingdom; Tumor and Microenvironment Interactions Group – Instituto de Investigacao e Inovacao em Saude, Universidade do Porto, Portugal; INEB-Instituto Nacional de Engenharia Biomédica, Portugal; Instituto de Patologia e Imunologia Molecular, Universidade do Porto, Portugal; Faculdade de Medicina, Universidade do Porto, Portugal; ICBAS-Instituto de Ciências Biomédicas Abel Salazar, Universidade do Porto, Portugal

**Author notes:** Medical Institute of Bioregulation, Kyushu University, Fukuoka, Japan (Takayuki Nojima).

## Abstract

Macrophages are essential cells of the immune system that alter their inflammatory profile depending on their microenvironment. Alternative polyadenylation in the 3’UTR (3’UTR-APA) and intronic polyadenylation (IPA) are mechanisms that modulate gene expression, in particular in cancer and activated immune cells. Yet, how polarization and colorectal cancer (CRC) cells microenvironment affect 3’UTR-APA and IPA in primary human macrophages remains unknown. Here, primary human monocytes were isolated from healthy donors, differentiated and polarized into a pro-inflammatory state and ChrRNA-Seq and 3’RNA-Seq were performed to quantify gene expression and characterize new 3’UTR-APA and IPA mRNA isoforms. Our results show that polarization of human macrophages from naïve to a pro-inflammatory state causes a marked increase both in proximal polyA site selection in the 3’UTR and in IPA events, in genes relevant for macrophage functions. Additionally, we found a negative correlation between differential gene expression and IPA during pro-inflammatory polarization of primary human macrophages. As macrophages are abundant immune cells in the CRC microenvironment that either promote or abrogate cancer progression, we investigated how indirect exposure to CRC cells affects macrophage gene expression and 3’UTR-APA and IPA mRNA events. Co-culture with CRC cells alters the inflammatory phenotype of macrophages, increases the expression of pro-tumoral genes and induce 3’UTR-APA alterations. Notably, some of these gene expression differences were also found in tumour-associated macrophages of CRC patients, indicating that they are physiological relevant. Upon macrophage pro-inflammatory polarization *SRSF12* is the pre-mRNA processing gene that is most upregulated. After *SRSF12* knockdown in M1 macrophages there is a global downregulation of gene expression, in particular in genes involved in gene expression regulation and in immune responses. Our results reveal new 3’UTR-APA and IPA mRNA isoforms produced during pro-inflammatory polarization of primary human macrophages and CRC co-culture that may be used in the future as diagnostic or therapeutic tools.

## Introduction

Communication between neighbouring cell types induces alterations in gene expression. In colorectal cancer (CRC), the worldwide second and deadliest form of cancer^1^, the recruited immune cells are influenced by the tumor heterogeneity and by their microenvironment^2^. Macrophages are abundant innate immune cells at the tumor microenvironment, that either cooperate with or abrogate cancer progression, depending on their inflammatory profile^3, 4^ and are involved in immune evasion, which is a hallmark of tumorigenic progression^5, 6^. Importantly, they show transcriptional programs that control pro-and anti-inflammatory cellular responses, enabling a spectrum of phenotypes to respond to each stimulus^7^. Naïve macrophages (hereby referred to as M0) may be polarized in a continuum of inflammatory populations, ranging from pro-inflammatory (M1-like, hereby referred to as M1) to anti-inflammatory (M2-like, hereby referred to as M2). At the onset of CRC progression, M1 macrophages recognize and attack tumor cells^8–10^. Yet, the cancer cells that escape, acquire increased invasive and metastatic properties, creating a tumor-permissive microenvironment, within which macrophages develop an profile similar to M2^4, 11^.

Transcriptomic studies have highlighted the number and biological relevance of mRNA isoforms produced by alternative polyadenylation in the 3’UTR (3’UTR-APA) and by intronic polyadenylation (IPA)^12^. Both APA and IPA contribute to gene expression regulatory mechanisms^13–15^, with consequences for cell proliferation, differentiation, and cell cycle progression^16–20^. APA occurs in more than 70% of human genes^21–24^, and determines mRNA half-life and subcellular location, as well as protein subcellular localization and function^25–27^. IPA occurs due to the recognition of polyadenylation signals (PASs) within introns, originating truncated transcripts that either are non-functional or code for proteins lacking the C-terminal^28^. IPA has been described in immune cells^29^, in leukaemia^30^ and occurs in physiological relevant genes such as in the core cleavage and polyadenylation machinery gene *PCF11*^31^. In cancer and activated T cells, mRNAs with short 3’UTRs due to 3’UTR-APA are more prevalent than those with long 3’ UTRs^15, 32–34^. It has also been shown that shorter IPA mRNA isoforms contribute to increased transcriptomic diversity in primary multiple myeloma cells and in naive B cells, memory B cells, germinal center B cells, CD5^+^ B cells, T cells and plasma cells^29^, but the prevalence of IPA events in primary human macrophages has not been reported so far.

Here, we address how polarization of primary human macrophages and co-culture with CRC cells affect fundamental processes such as gene expression, 3’UTR-APA and IPA, in order to identify transcriptomic alterations in relevant mRNA isoforms, that we named signatures. We isolated primary human monocytes from healthy donors, exposed them to pro-inflammatory conditions and subsequently co-cultured them with two different CRC cell lines. By 3’ RNA-Seq, we identified a robust 35-gene signature of differentially-expressed genes (DEGs) in M1 polarized macrophages, comprising of inflammatory and immune-related genes. Noteworthy, some of these gene expression alterations are observed in tumor-associated macrophages in CRC patients. In addition, M1 polarization of primary human macrophages induces proximal PAS usage, leading to the production of mRNAs with short 3’UTRs in physiologically relevant genes, such as *IL17RA* and *TP53RK*. IPA events are increased in M1 polarized macrophages affecting genes such as *MAP3K8* and *WRD33*. Both 3’UTR-APA shortening and IPA are further upregulated after CRC co-culture, in particular for *MAP3K8* and *WRD33*. *SRSF12* expression is strongly upregulated in primary human macrophages upon M1 polarization. Notably, *SRSF12* knockdown causes a strong downregulation in the expression of genes involved in regulation of gene expression and in macrophage functions. Moreover, *SRSF12* knockdown leads to an increase in *MBNL1* proximal PAS selection and an increase in *CDC42* IPA. Our results describe 3’UTR-APA and IPA events in primary human macrophages driven by M1 polarization and by exposure to colorectal cancer cells, which are relevant for gene expression regulation and macrophage functions. Furthermore, our results a new function for *SRSF12* in pro-inflammatory macrophages, key cells in tumour response.

## Results

### Pro-inflammatory polarization alters primary human macrophages transcription profile

Macrophages can be experimentally differentiated from quasi-naïve, differentiated and unpolarized cells into a spectrum of inflammatory profiles. We obtained primary human CD14^+^ monocytes isolated from peripheral blood mononuclear cells (PBMCs) of healthy blood donor buffy coats, differentiated them into macrophages in the presence of M-CSF (herein referred as M0) and polarized them towards pro-inflammatory conditions (M1) by incubation with LPS and IFN-γ, following a well-established methodology^35–37^ represented in Figure 1A. M0 macrophages are predominantly round/ameboid-like, while M1 macrophages present a fibroblast-like morphology (Figure 1A). Characterization of the macrophage inflammatory profile shows that upon M1 polarization the CD14 monocytic cell lineage marker is maintained (Figure S1A), the expression of CD86, CCR7 and IL1β pro-inflammatory markers is increased, and the expression of CD163 and TGF-β anti-inflammatory markers is decreased (Figures S1B-1E), thus showing that M1-polarized macrophages present a *bona fide* pro-inflammatory phenotype.

**Figure 1.**
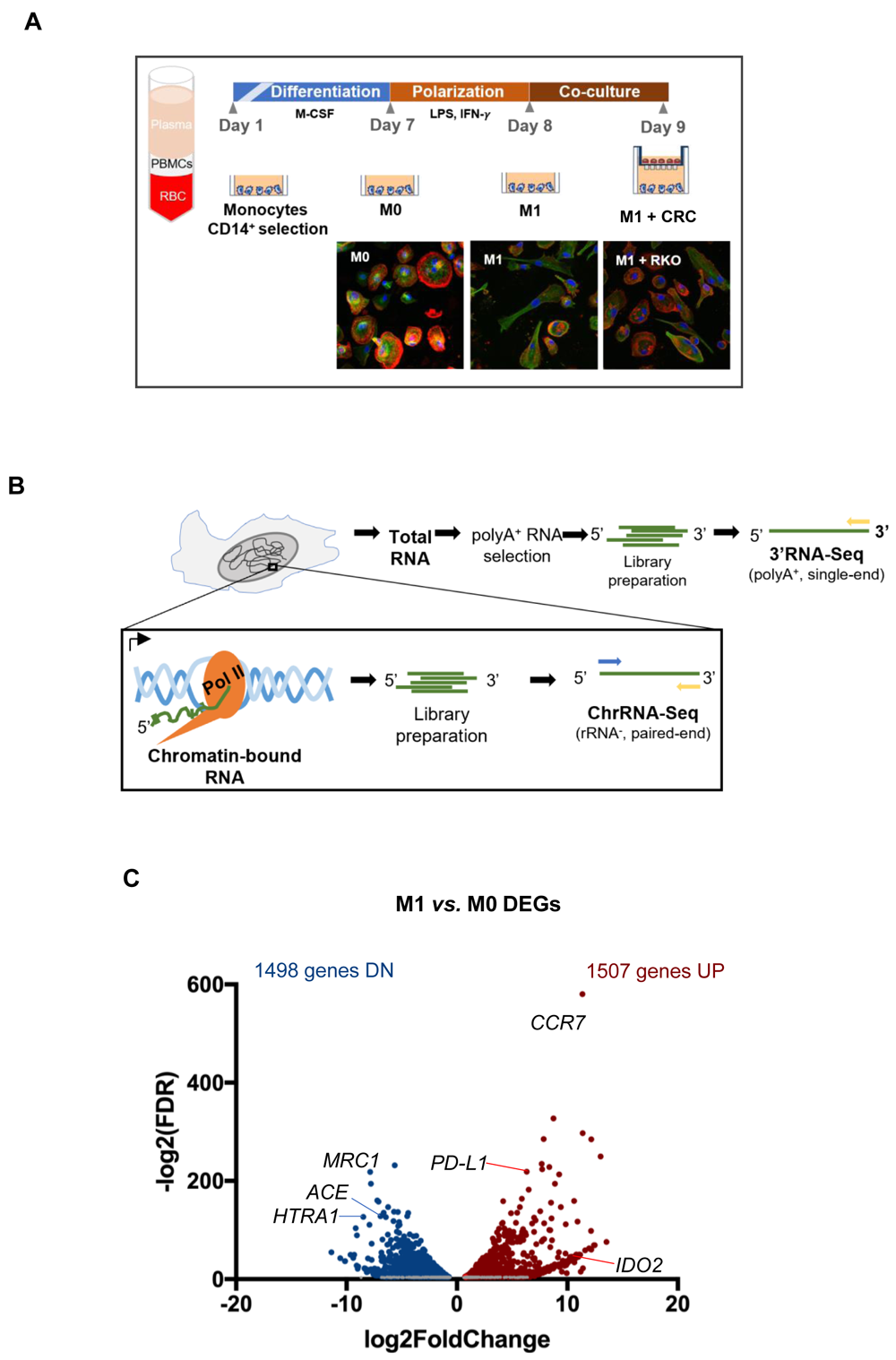
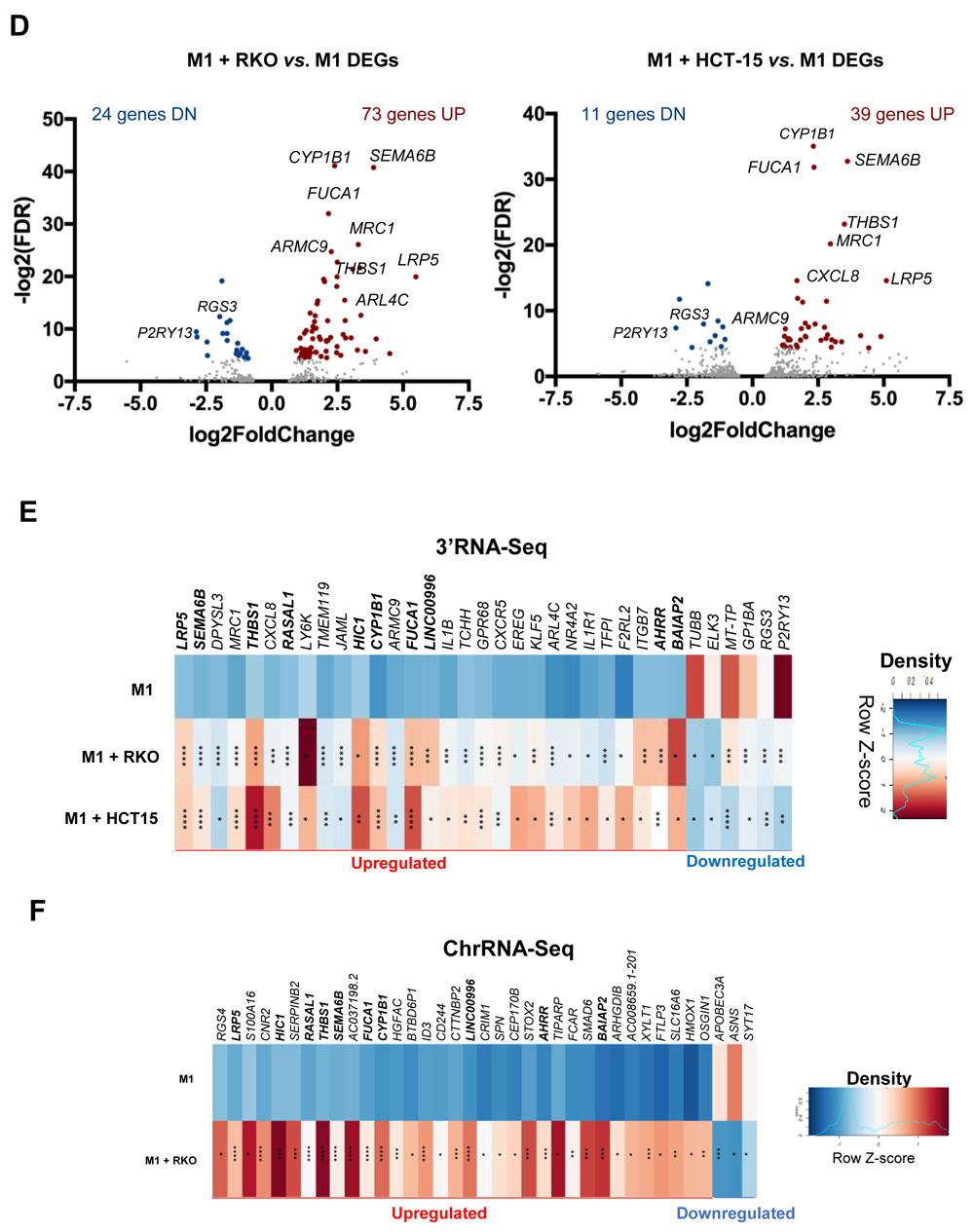
M1 polarization and CRC co-culture modulate macrophage inflammatory and gene expression profiles. **A.** Top, experimental setup: PBMCs are isolated through gradient centrifugation and then exposed to CD14+ beads to select CD14+ monocytes. Monocytes are differentiated for 7 days to obtain unpolarized naïve M0 macrophages and then polarized to M1-like (M1) macrophages for 24 hours. M1 macrophages are co-cultured with CRC cell lines RKO or HCT15 for 24h. Bottom, confocal microscopy images show actin-tubulin cytoskeleton of (**M0**), LPS+IFNγ stimulated pro-inflammatory (**M1**) polarized macrophages, and LPS+IFNγ-stimulated macrophages co-cultured with CRC cell line (**M1 + RKO**). DNA stained in blue, actin stained in red, tubulin stained in green. **B.** Setup of extraction and library preparation for RNA-Seq of primary human macrophages. rRNA-depleted chromatin-bound RNA were converted into paired-read libraries for ChrRNA-Seq, and polyA^+^-enriched total RNA were converted into single-end libraries for 3’RNA-Seq. **C**. Volcano plot of 3’RNA-Seq data comparing differentially expressed genes (DEGs) in M1 *vs*. M0. Blue and red dots represent statistically significant genes with log2(Fold Change) <-1 or >1, respectively. **D**. Volcano plot of 3’RNA-Seq data comparing differentially expressed genes (DEGs) in M1 macrophages co-cultured with RKO cells (M1+ RKO, left) or HCT-15 cells (M1 + HCT15, right) *vs*. M1 macrophages in monocultures (M1). Red dots represent statistically significant upregulated genes (log2(Fold Change >1); blue dots represent statistically significant downregulated genes (log2(Fold Change <-1). Grey dots represent non-significantly expressed genes. **E**. Representative donor heatmap of 3’RNA-Seq DEGs in macrophages co-cultured with RKO (M1 + RKO) or HCT15 (M1 + HCT15) *vs*. M1 macrophages in monocultures, ordered by fold change. n=3 healthy donors, p-value calculated using DESeq2 test through the Benjamini-Hochberg method: *=p< 0,05; **= p <0,01; ***=p< 0,005; **** = p<0,0001. **F.** Representative donor heatmap of ChrRNA-Seq DEGs in macrophages co-cultured with RKO (M1 + RKO) *vs*. M1 macrophages, ordered by fold change. n=3 healthy donors, ChrRNA-Seq data p-value calculated using DESeq2 test through the Benjamini-Hochberg method: *=p< 0,05; **=p<0,01; ***=p<0,005; ****=p<0,0001. Genes in bold are upregulated in 3’RNA-Seq and ChrRNA-Seq data.

To assess alterations in differential gene expression, 3’UTR-APA and IPA events upon macrophage polarization, we performed 3’RNA-Seq, which provides both quantification of gene expression based on RNA levels at the 3’ end and APA profiles^36^ and also chromatin-bound RNA-Seq (ChrRNA-Seq) to pinpoint those events that occur in nascent RNA^38^ (Figure 1B). 3’ RNA-Seq results reveal that 3005 genes show a significantly altered expression in M1 in comparison to M0 macrophages (Figure 1C). The upregulated 1507 gene subset was significantly enriched in Gene Set Enrichment Analyses (GSEA) terms related to stress and immune responses, IFN-γ and TNF-α signalling pathways, all of which are characteristic of pro-inflammatory responses (Figure S1F). Of note, the pro-inflammatory markers *CCR7*, *CCL19*, *IDO2*, *IL7R* and the *PD-L1* immune checkpoint gene are present among the upregulated gene subset (Figure 1C and Supplementary Data 1). Conversely, the downregulated 1498 gene expression subset (Figure 1C and Supplementary Data 1) was significantly enriched in GSEA terms associated with extracellular matrix (ECM), secretory vesicles and endocytosis, lipid binding, and GPCR signalling (Figure S1F), which are classically associated with anti-inflammatory macrophages^39^.These results indicate that M1 polarization leads to a phenotypically and transcriptomically robust pro-inflammatory profile.

### CRC co-culture induces a 35-gene expression signature in primary human macrophages

At the early stages of CRC progression, tumor-associated macrophages (TAMs) are polarized towards M1, a tumor-inhibitory phenotype^8–10^. Therefore, we polarized macrophages into M1 and co-cultured them with the CRC cell lines RKO and HCT-15^40^ to analyse the effect of cancer cells exposure on the macrophage transcriptome (Figures 1A). Both CRC cell lines belong to CMS1 (consensus molecular subtype 1), characterized by hypermutation, high microsatellite instability and pronounced immunogenicity^41^, but differ in several tumorigenic markers such as *APC*, *K*-*ras*, *B-raf*, *TGFBR2* and *MLH1*^42^. By using these two CRC cell lines we can ascertain a specific effect to an inherent characteristic of each cell line, or determine if the effect could be due to a more general feature of CRC exposure. Upon co-culture with CRC cells, macrophage morphology reverts to an ameboid-like phenotype, similar to M0, a phenotypic alteration constant in all replicates and with both CRC cell lines (Figure 1A, S2A), whilst still maintaining the CD14 and CD86 markers (Figure S2B-C). 3’RNA-Seq analyses of the macrophages revealed that RKO and HCT-15 co-culture induces robust alterations in gene expression (Figure 1D and Supplementary Data 1) and the 3’RNA-Seq Pearson correlation analysis indicate a very high correlation between biological replicates (Figure S2D).

We identified 35 differentially expressed genes (DEGs) in macrophages that are common to both CRC cell lines co-culture, including 29 upregulated and 6 downregulated genes (Figures 1E). The 35 DEGs signature includes 15 anti-inflammatory and/or pro-tumorigenic genes, including *LRP5, SEMA6B*, *MRC1, ARMC9, FUCA1, EREG* and *ARL4C*, genes related to immune modulation and inflammation, such as *THBS1, CXCL8*, *LY6K* and *KLF5*, and pro-inflammatory genes, such as *IL1B, CXCR5, IL1R1* and *F2RL2*, and some of these DEGs were validated by RT-qPCR (Figure S2E). Other anti-inflammatory and/or pro-tumoral genes significantly upregulated DEGs in macrophages co-cultured with CRC cells include *SERPINB2*, *MS4A6A*, *TGFA*, *FOSB*, *FABP4* and *RGS1* (Figure S2F). Upregulated genes in the CRC-exposed macrophage signature have GSEA terms associated to vesicle-mediated transport and endocytosis, to response to IL-1-mediated transport, leukocyte chemotaxis and migration (Figure S2G), which are associated with macrophage recruitment to tumors and with tumor progression41,43-46. Some of the 35 DEGs are included in the chromatin-bound RNA of co-cultured macrophages (Figure 1F, Supplementary Data 1). ChrRNA-Seq analyses show a slight increase in global transcription in co-cultured macrophages with CRC cells (Figure S3A). Overall, genes with differential expression obtained by ChrRNA-Seq in co-cultured macrophages shared many of the GSEA terms associated to 3’RNA-Seq, such as regulation of cell differentiation, lipid metabolism and the Wnt pathway (Figure S3B).

Taken together, these results indicate that macrophages change their inflammatory profile and adopt pro-tumorigenic characteristics upon CRC co-culture.

### CRC patients show the same gene expression alterations as those present in the 35-gene signature

Some of the most upregulated genes found in CRC co-cultured macrophages – *LRP5, DPYSL3*, *THBS1, CXCL8*, *IL1ß* and *EREG* (Figure 1E) – are related to the Wnt pathway^45, 47–51^, which is constitutively activated in CRC^52^. *LRP5* is the most upregulated gene in macrophages that were co-cultured with CRC cells as shown by the heat map, the Integrative Genomics Viewer (IGV) of the ChrRNA-Seq and 3’RNA-Seq data (Figure 2A) and by the 2-fold increase in LRP5 protein levels observed by western blot (Figure 2B). To evaluate whether this upregulation occurs in clinically-relevant contexts, we analysed gene expression levels in CRC patients, using RNA-Seq data from The Cancer Genome Atlas (TCGA) database and selecting CD68^High^ tumor associated macrophages (TAMs). Notably, *LRP5* is upregulated in macrophages from CRC patients, which correlates with an increase in patient survival (Figure 2C). Furthermore, several other genes in the 35 gene DGE signature presented by macrophages after co-culture with CRC (from Figure 1E), including *EREG, CXCL8*, *IL1B, ARL4C*, *ARMC9*, *BAIAP2*, *F2RL2*, *GP1BA*, *MT-TP* and *P2RY13*, show the same expression profile in TAMs from CRC patients (Figure 2D). These results indicate that CRC co-culture induces the expression of genes that are similarly expressed by TAMs from CRC patients, which are physiologically relevant for tumor response.

**Figure 2.**
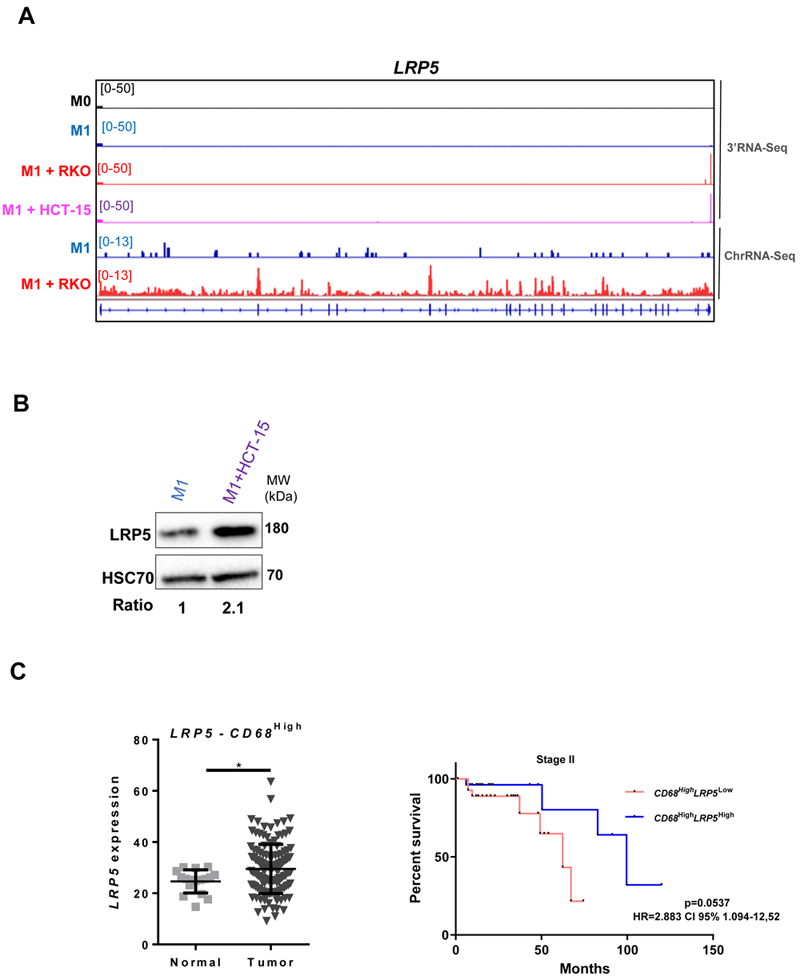
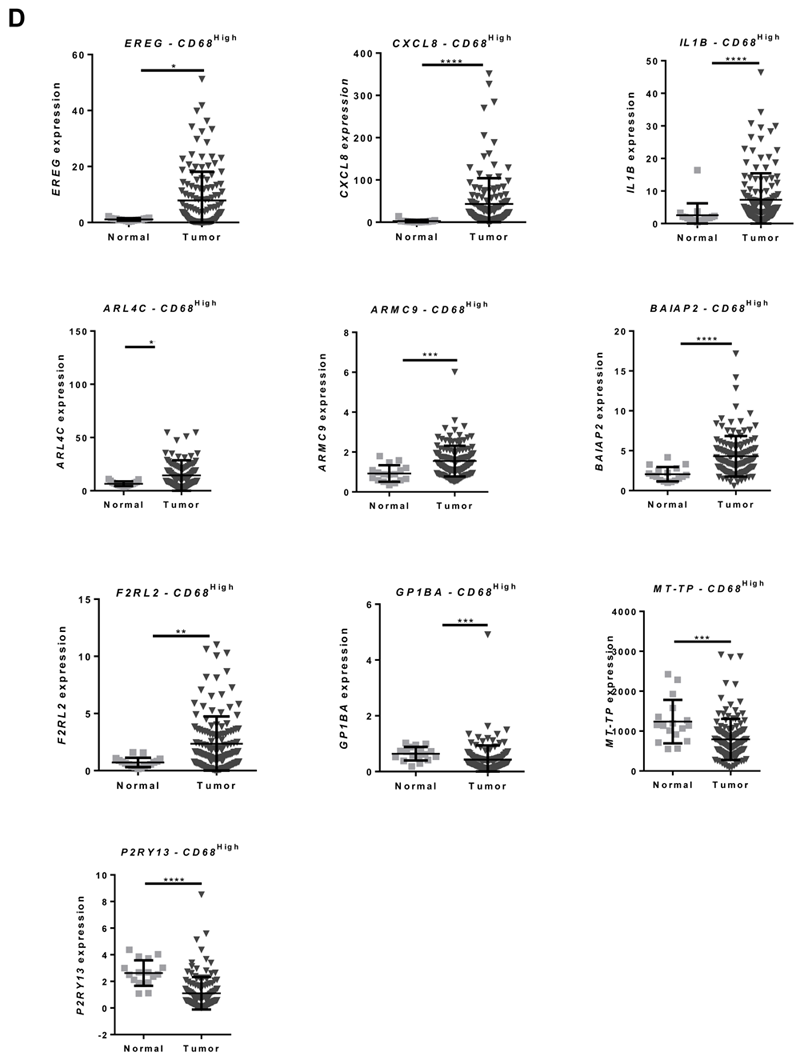
C patients present differential expression of inflammation-related genes that is also found in CRC co-cultured macrophages. **A.** *LRP5* representative gene profile and transcript counts in 3’RNA-Seq (top) and ChrRNA-Seq (bottom). M0 macrophages (black), M1 macrophages cultured alone (blue) and M1 macrophages co-cultured with the CRC cell lines RKO (red) or HCT-15 (lilac) visualized in IGV. B. Western blot showing that LRP5 protein levels also increase upon CRC co-culture. C. Expression data from CRC patients from the TCGA database: microarray and RNA-Seq gene expression of *LRP5* in CD68^High^ macrophage population between normal and tumor tissue; Right, Kaplan-Meier survival curves of CD68^High^LRP5^Low/High^ patients in tumour stage II. D. *EREG*, *CXCL8*, *IL1B ARL4C*, *ARMC9*, *BAIAP2*, *F2RL2*, *GP1BA*, *MT-TP* and *P2RY13* expression in CRC patients in CD68^High^ macrophage population between normal and tumor tissue, data from TCGA database.

### Pro-inflammatory polarization induces 3’UTR-APA shortening and IPA in primary human macrophages

To investigate the effect of M1 polarization in 3’UTR-APA and IPA we analysed the 3’RNA-Seq data of M1 *vs*. M0 macrophages, using APAlyzer^53^ and obtained a dataset consisting of APA and IPA mRNA isoforms relative expression differences (Supplementary Data 2).

We observed a strong upregulation of mRNA isoforms with short 3’UTRs resultant from proximal PAS selection as 141 genes showed upregulation of short 3’UTRs compared to 6 genes showing upregulation of long 3’UTRs (Figure 3A). Genes that undergo 3’UTR shortening showed GO terms involved in regulation of gene expression, and also related to the immune system, cell-cell communication, extracellular exosomes and intracellular transport (Figure 3B), which are important for macrophage inflammatory functions. *IL17RA* (Interleukin 17 receptor A) is illustrative of a gene that shows upregulation of the shortest 3’UTR mRNA isoform and downregulation of the longest 3’UTR in M1 *vs*. M0 (Figure 3C). Interleukin 17A and its receptor IL17RA play a pathogenic role in many inflammatory and autoimmune diseases and IL17RA contributes to the inflammatory response by inducing recruitment of innate immune cells^54^. In addition, IL17RA deletion predicts high-grade colorectal cancer and poor clinical outcomes^55^. *TP53RK* (TP53 Regulating Kinase) is another example of a gene that displays a decrease in the expression of the longest 3’UTR mRNA isoform and an upregulation of the shortest 3’UTR mRNA isoform in M1 *vs* M0 (Figure 3D). TP53RK/PRPK enables p53 binding activity and its kinase activity, and the expression levels of phosphorylated TP53RK/PRPK are higher in metastatic CRC tissues in comparison to normal tissues^56^. Other physiologically relevant genes with upregulated short 3’UTRs upon M1 polarization include *YKT6*, which is connected to the Wnt pathway, lysosome fusion, and exosome secretion^57^, *ALYREF*, an RBP that is recruited to target mRNAs through interaction with IRAK2 and binds 5’ and 3’ UTRs through a complex with CSTF2/CstF64^58, 59^ and *MYO10*, whose 3’UTR is regulated by TGFβ signalling via SMAD transcription factors^60^ (Supplementary Data 2). Of note, the 3’UTR-APA changes induced by M1 polarization do not present a significant correlation with differential gene expression, as shown by the two-tailed Pearson correlation between significant genes with both DGE and isoform RED in differential APA in M1 *vs* M0 (Figure 3D).

**Figure 3.**
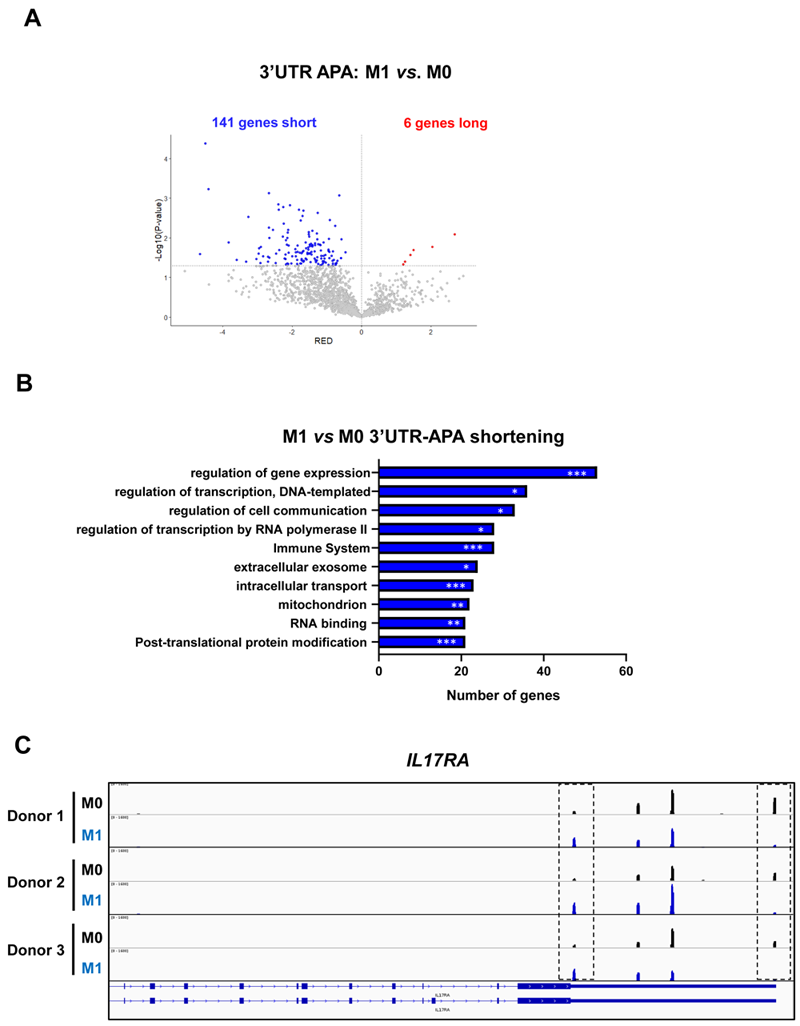
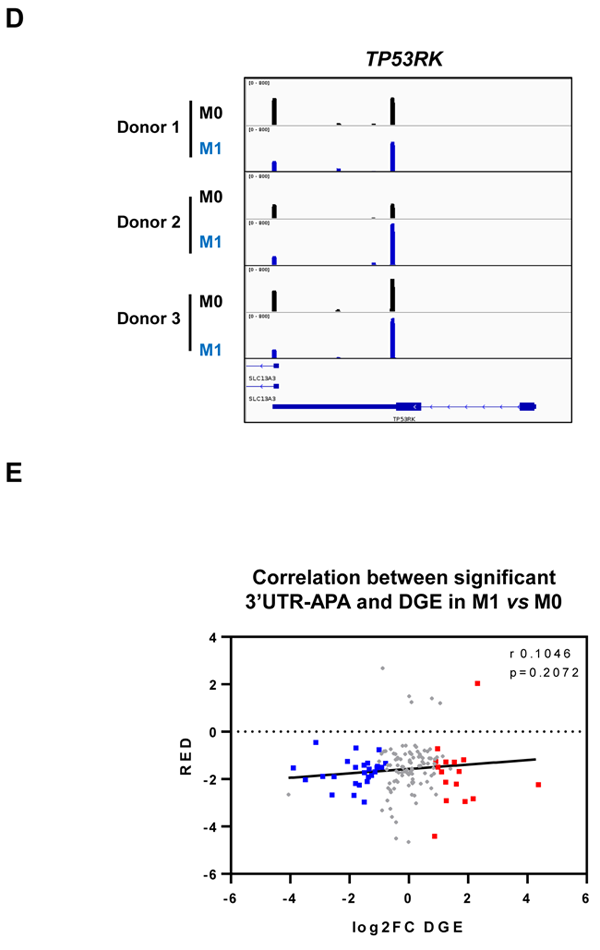
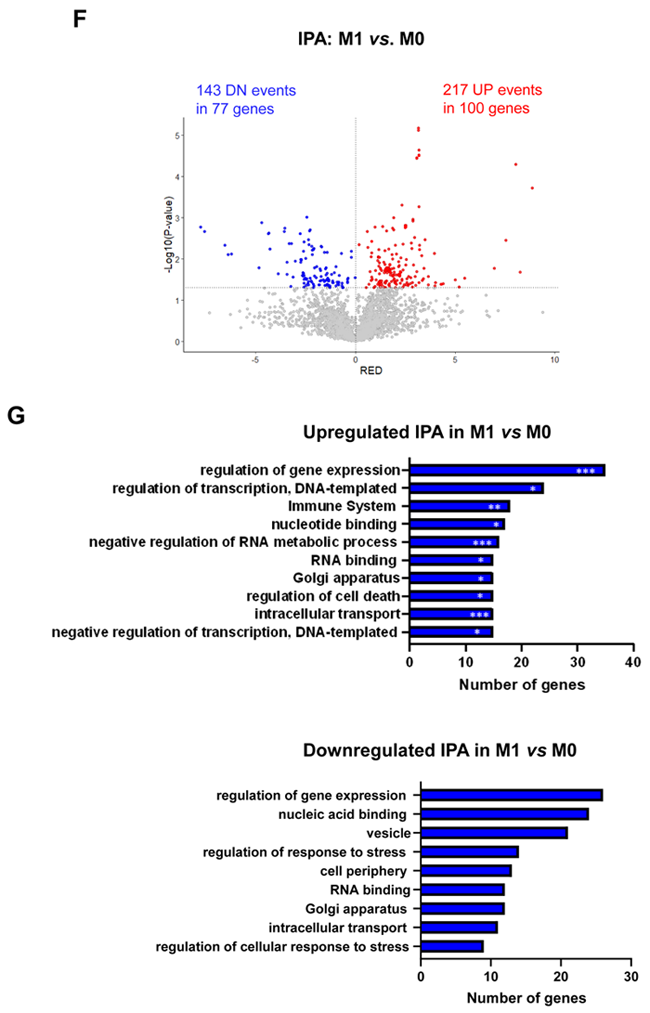
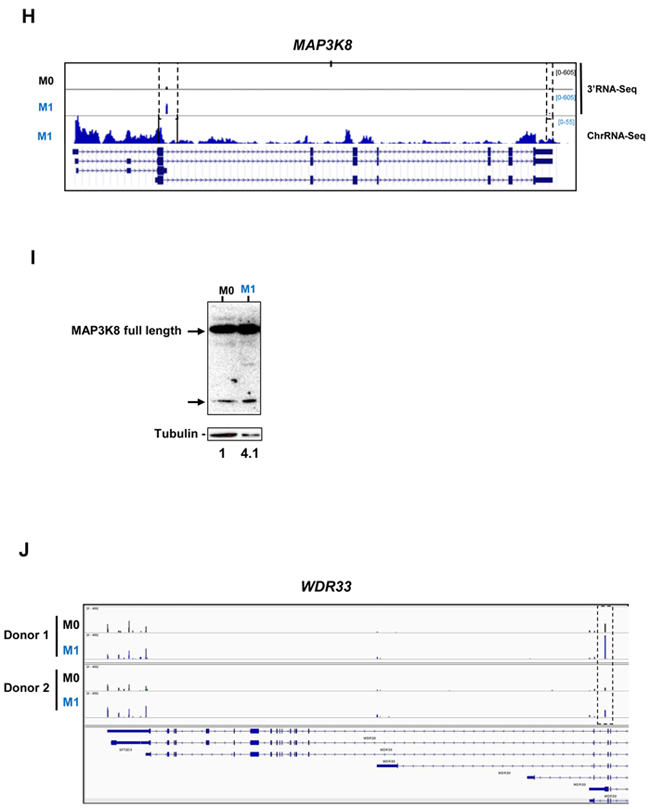
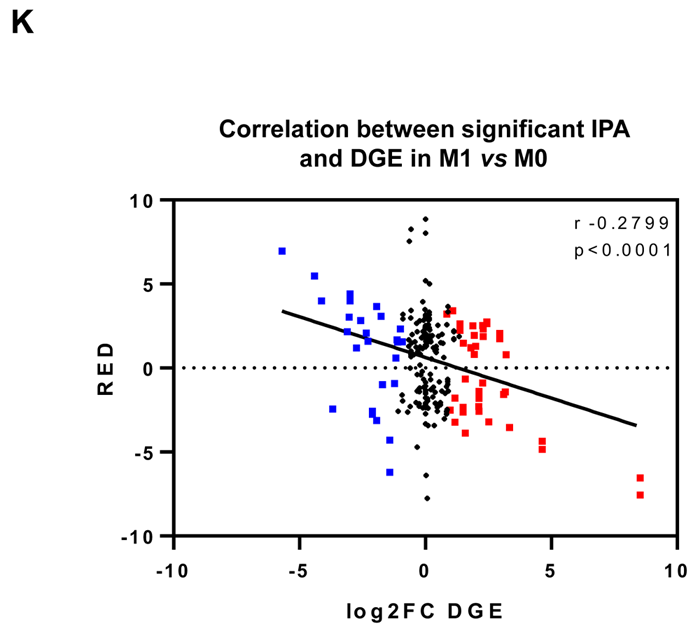
M1 polarization induces 3’UTR-APA shortening and intronic polyadenylation (IPA). **A.** Volcano plot of genes showing differential 3’UTR-APA mRNA isoforms in M1 *vs*. M0. Red dots represent genes with genes with Relative Expression Differences (RED)>0, corresponding to expression of the long isoform, and blue dots represent genes with RED<0, corresponding to expression of the short isoform and p<0.05, grey dots represent genes without significantly different RED. **B**. Top 10 Gene Ontology terms in M1 *vs*. M0 of genes showing upregulated short 3’UTR. Gene Ontology p-value calculated using Fisher’s test (Panther). *=p<0,05; **=p<0,01; ***=p<0,005. **C.** *IL17RA* gene profile and transcript counts in 3’RNA-Seq visualized in IGV for 3 different healthy donors. M0 macrophages (black), M1 macrophages cultured alone (blue) visualized in IGV. Dashed boxes indicate the reads correspondent to short (proximal PAS usage) and long (distal PAS usage) 3’UTRs mRNA isoforms. **D.** *TP53RK* gene profile and transcript counts in 3’RNA-Seq visualized in IGV for 3 different healthy donors. M0 macrophages (black), M1 macrophages cultured alone (blue) visualized in IGV. **E.** Two-tailed Pearson correlation between significant differential 3’UTR-APA genes (RED) and DGE (log2FC DGE) in M1 *vs* M0 macrophages (n=3 donors, r 0.1046, p=0.2072). Genes shown in blue have significant downregulation of expression, genes shown in red have significant upregulation of expression. **F.** Volcano plot of genes showing differential IPA mRNA isoforms in M1 *vs*. M0 analysed by Apalyzer (genes that undergo an increase in IPA events are represented in red (RED>0), UP: upregulated IPA; genes that undergo a decrease in IPA events, hence upregulating expression of last exon are represented in blue, DN: downregulated IPA, RED < 0) and p < 0.05. **G.** Top 10 Gene Ontology terms in M1 *vs*. M0 of genes showing differential IPA. Gene Ontology p-value calculated using Fisher’s test (Panther). *=p<0,05; **=p<0,01; ***=p<0,005. **H**. Gene profiles and transcript counts of *MAP3K8* obtained through 3’-RNA-Seq (**top**) and ChrRNA-Seq (**down**) of M0 macrophages (**black**) and M1 macrophages (**blue**), visualized in IGV. **I**. *MAP3K8* protein levels measured by western blot using tubulin as loading control. Arrows point to full-length MAP3K8 and to an isoform of smaller molecular weight. Numbers below the gel correspond to the ratios of the smaller protein band in M1 *vs* M0, quantified by densitometry. **J.** IGV gene profiles and transcript counts of *WDR33* obtained through 3’RNA-Seq of M0 macrophages (**black**) and M1 macrophages (blue) in two different donors. Dashed box indicates the reads correspondent to the IPA mRNA isoform. **K.** Two-tailed Pearson correlation between significant differential IPA (RED) and DGE (log2FC DGE) in M1 *vs* M0 macrophages (n=3 donors, r -0.2799, p<0.0001). Genes shown in blue have significant downregulation of expression, genes shown in red have significant upregulation of expression.

When we analysed IPA, we observed a strong increase in the expression of IPA mRNA isoforms in M1 *vs* M0, as 100 genes showed upregulation of IPA mRNA isoforms, in a total of 217 upregulated IPA events (Figure 3F). Upregulation of IPA isoforms showed GO terms connected to regulation of gene expression and the immune system (Figure 3G), similarly to the GO terms found for 3’UTR-APA events, while downregulation of IPA mRNA isoforms present GO terms that curiously do not include the immune system, but include regulation of response to stress (Figure 3G). Genes that undergo IPA upregulation upon M1 polarization include *MAP3K8*, which is widely expressed in immune cells and tumors^61^. In M0, *MAP3K8* uses two PAS, one at the 3’UTR and another PAS in an intron. In M1 the intronic PAS is selected ∼3-fold more efficiently than in M0 (Figure 3H). This shorter transcript is described in ENSEMBL and corresponds to a 132 aa, 15kDa MAP3K8 truncated protein isoform predicted by the Uniprot database, Q5T854. Interestingly, by western blot we observe an increase in the intensity of a protein band of molecular weight similar to the predicted Q5T854 protein (Figure 3I). It remains to be investigated whether this protein corresponds to the IPA mRNA isoform and if it is functional. *WDR33*, which codes for the protein that binds to the PAS^62^, is another gene that undergoes a >2-fold increase in an IPA mRNA isoform during pro-inflammatory polarization (dashed bow in Figure 3J). This IPA isoform, if functional, produces a shorter WDR33 protein of still unknown function.

Interestingly, we found a strong negative correlation between IPA and gene expression as shown by the two-tailed Pearson correlation between significant genes with both DGE and isoform RED in differential IPA in M1 *vs* M0 macrophages (Figure 3K). Genes with an increase in IPA tend to be downregulated, and genes with a decrease in IPA tend to be upregulated.

### CRC co-culture affects 3’UTR-APA and IPA in primary human macrophages

We next analysed how co-culture with CRC cells affects 3’UTR-APA and IPA in primary human macrophage by analysing 3’RNA-Seq data with APAlyzer (Supplementary Data 2).

Co-culture of M1 primary human macrophages with both RKO and HCT-15 CRC cell lines induces 3’UTR-APA changes, with an increase in the expression of mRNA isoforms with short 3’UTRs (Figures 4A and 4B). The list of genes with differential 3’UTR-APA that are common to both co-cultured macrophages datasets are listed in Table I. Interestingly, common upregulated long 3’UTR genes include *C11orf54* and *STARD3NL*, which are involved in exosome functions^63, 64^ and *TMEM265*, whose expression is negatively correlated with tumor-infiltrating immune cells^65^, while upregulated short 3’UTR genes include *PBX3* (Figure 4C), which is associated with inflammation and promotes migration and invasion of colorectal cancer cells^66–68^.

**Figure 4.**
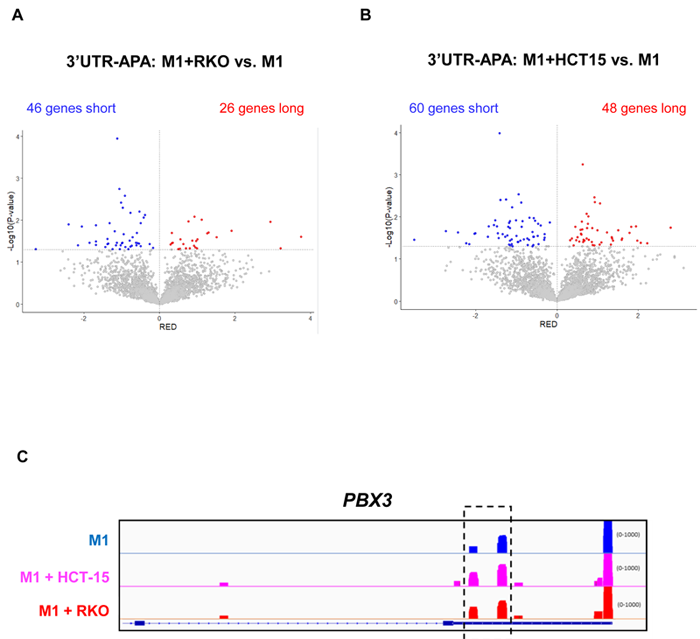
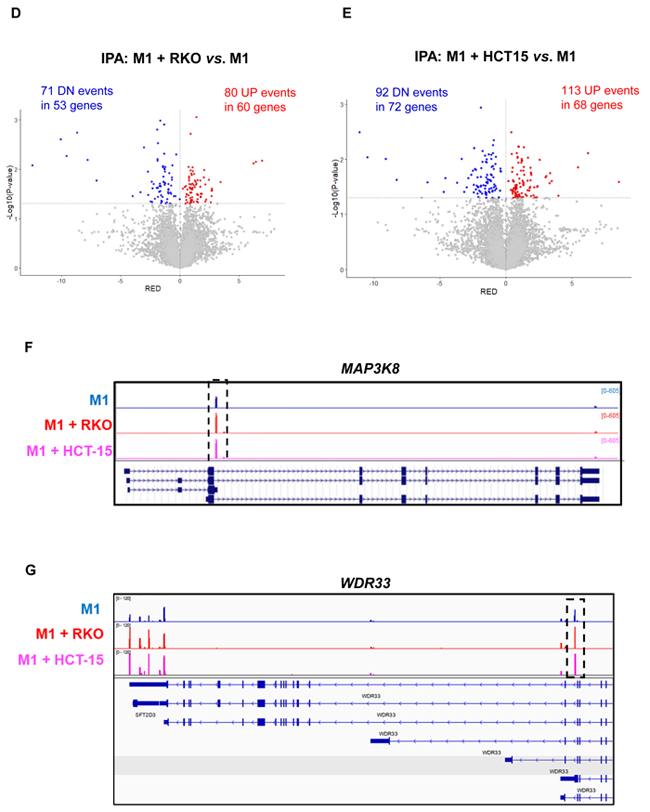
CRC co-culture induces 3’UTR-APA and IPA changes in macrophages. **A-B.** Volcano plots of genes showing differential 3’UTR-APA changes in M1+RKO *vs*. M1 (A) and in M1+HCT15 *vs*. M1 (B). Genes that express short 3’UTRs mRNAs are represented in blue and genes that express long 3’UTRs mRNAs are represented in red. **C**. IGV gene profiles and transcript counts of *PBX3* obtained through 3’RNA-Seq of M1 macrophages (**blue**) and M1 macrophages co-cultured with the CRC cell lines HCT-15 (**lilac**) and RKO (**red**). Dashed box indicates an increase in reads over proximal polyA sites, in CRC co-cultured macrophages. **D-E.** Volcano plots of genes showing IPA changes in M1+RKO *vs*. M1 (D) and in M1+HCT15 *vs*. M1 (E). Genes that express more IPA mRNAs are represented in red (UP) and genes that express less IPA mRNAs are represented in blue (DN). **F**. IGV gene profiles and transcript counts of *MAP3K8* obtained by 3’RNA-Seq of M1 macrophages (**blue**) and M1 macrophages co-cultured with the CRC cell lines RKO (**red**) and HCT-15 (**lilac**). Dashed box indicates an increase in IPA reads in CRC co-cultured macrophages. **G.** IGV gene profiles and transcript counts of *WDR33* obtained by 3’RNA-Seq of M1 macrophages (blue) and M1 macrophages co-cultured with the CRC cell lines RKO (**red**) and HCT-15 (**lilac**). Dashed box indicates an increase in IPA reads in CRC co-cultured macrophages.

**Table I.** Differential 3’UTR-APA genes in M1+CRC vs M1. Alphabetical lists of genes showing differential 3’UTR-APA mRNA isoforms that are common to the two macrophage datasets correspondent to the co-cultures with RKO and HCT-15 cell lines.

| Short 3'UTR | Long 3'UTR |
| --- | --- |
| *GGH* | *C11orf54* |
| *KDM7A* | *DHDDS* |
| *LMAN1* | *NDOR1* |
| *PBX3* | *STARD3NL* |
| *RBMX* | *TIMM9* |
| *SPCS2* | *TMEM265* |

We also found that, after co-culture with CRC cells, macrophages display an increase in IPA (Figure 4D and 4E). The genes that are common to both co-cultured macrophages datasets in the IPA analysis include 17 genes with upregulated IPA (Table II). This list comprise physiologically relevant genes for macrophage and cancer biology, such as *MAP3K8* (Figure 4F). We had initially observed that pro-inflammatory polarization caused a ∼2-fold increase in a *MAP3K8* IPA mRNA isoform (Figure 3H). Notably, co-culture with CRC cells further increases by ∼2-fold the expression of this IPA mRNA isoform (Figure 4F). *WDR33*, is another interesting example of a gene that expresses an IPA isoform which increases by pro-inflammatory polarization (Figure 3H) and that is further increased in macrophages after co-culture with CRC (Figure 4G). Curiously, the bioinformatic analysis included this gene in the downregulated IPA common list of genes (Table II), because the 3’UTR-APA long mRNA isoform also increases. Other genes with upregulated IPA are *CUX1*, which inhibits NF-kB transcriptional activity and contributes toward tumor progression and is a target of TGFß^69^ and *MESD*, a chaperone necessary for LRP5 translocation, which is related to the Wnt pathway^70, 71^ and *TIA1*, a RBP expressed in activated macrophages and in CRC^72,73^.

**Table II.**
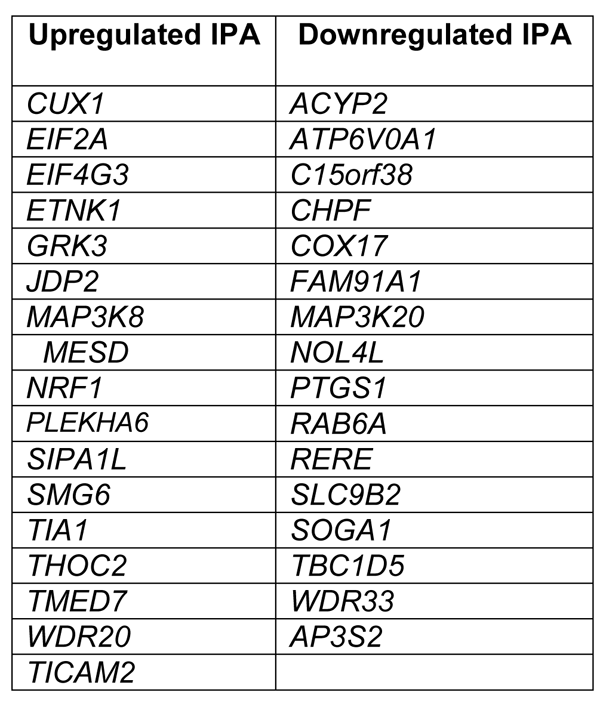
Differential IPA genes in M1+CRC vs M1. Alphabetical lists of genes showing upregulated and downregulated IPA that are common to the two macrophage datasets correspondent to the co-cultures with RKO and HCT-15 cell lines.

The fifteen genes common to both co-cultured macrophages datasets that show downregulated IPA also have a physiological function in macrophage and cancer biology (Table II). This list include *COX17*, which provide essential copper transport for M1 macrophages in tumors^74^, *RAB6A*, associated with Golgi regulation of TNF secretion in macrophages^75^ and *SLC9B2*, which is associated with infiltrating macrophages in CRC^76^.

Curiously, some of the genes in the APA and IPA lists are connected to the Wnt pathway and most of them are related to lipids metabolism, through functions related to the Golgi, ER, mitochondria, or plasma membranes, and exosomes.

Our results show that polarization of primary human macrophages leads to an increase in the expression of 3’UTR-APA short mRNA isoforms and in IPA mRNA isoforms, in physiologically relevant genes. Moreover, CRC co-culture induces even further APA and IPA changes in macrophages.

### *SRSF12* knockdown decreases gene expression in M1 macrophages

To understand the mechanisms behind the transcriptomic alterations observed following macrophage M1 polarization, we analyzed the 3’RNA-Seq data to identify changes in the expression of genes that code for protein factors involved in cleavage and polyadenylation, splicing and transcription termination (Figure 5A). Of these genes, the most upregulated gene in M1 macrophages is the splicing repressor *SRSF12/SRrp35* ^77^ (Figure 5A and B). Therefore, we knocked down *SRSF12* from primary macrophages after M1 polarization using siRNAs (Figure 5C) and achieved >80% of *SRSF12* mRNA depletion (Figure 5D). Interestingly, *SRSF12*-depleted macrophages show a marked downregulation of gene expression (333 downregulated genes *vs.* 14 upregulated genes) (Figure 5E and Supplementary Data 3). Notably, several genes involved in gene expression regulation are downregulated, in particular *CSTF2/CstF64, SRSF2*, *SRSF3*, *U2SURP*, *DBR1*, *PARP2*, *EIF5A2*, as well as several transcription factors such as *NFYA*, *EGR1*, *ELK3*, *SMAD1* and *MTA3* (Figure 5F, Supplementary Data 3). Other downregulated genes are involved in macrophage-relevant functions, such as regulation of immune responses and leukocyte proliferation (Figure 5F).

**Figure 5.**
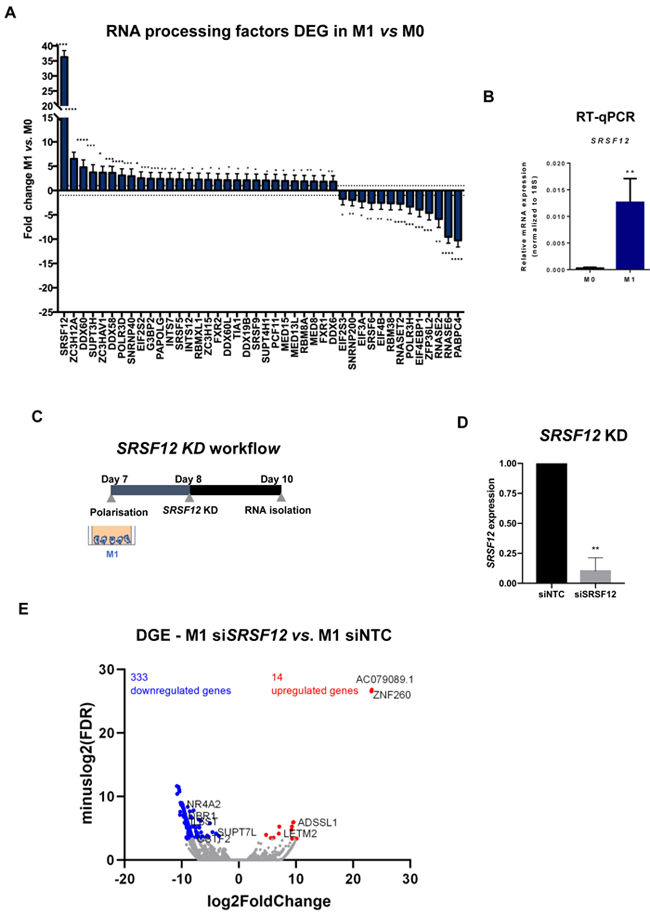
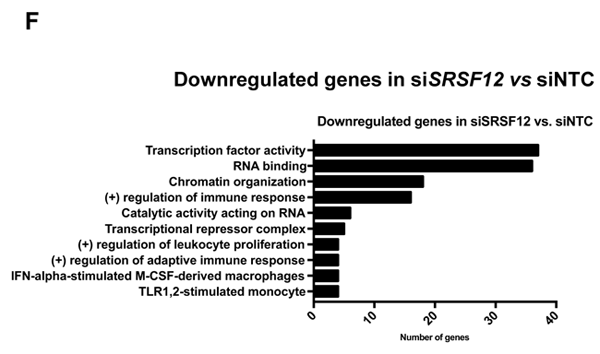
SRSF12 modulates gene expression in pro-inflammatory macrophages. **A.** Fold change distribution of CPA and transcription regulation DEGs in M1 vs. M0, ordered by fold change. n=3 healthy donors, p-value calculated using DESeq2 test through the Benjamini-Hochberg method: *=p< 0,05; **= p <0,01; ***=p< 0,005; **** = p<0,0001. **B.** *SRSF12* mRNA levels in M1 *vs* M0 quantified by RT-qPCR. n=4 donors, Student’s t-test, p = 0.0202. **C.** Experimental setup of *SRSF12* knockdown in primary human macrophages. **D**. Levels of *SRSF12* knockdown quantified by RT-qPCR. n=3 donors, Student’s t-test, p = 0.0050 **E.** Volcano plot of DEGs in si*SRSF12 vs* siNTC in M1 macrophages in n=2 donors. Red dots represent genes with log2(Fold Change) >1, blue dots genes with log2(Fold Change) <– 1 and p < 0.1, grey dots represent non-significantly expressed genes. **F**. GSEA terms of downregulated DEGs in si*SRSF12 vs* siNTC. n= 2, p-value calculated using GSEA: * = p < 0,05; ** = p < 0,01; *** = p < 0,005.

We next analyzed 3’UTR-APA and IPA events in si*SRSF12 vs* siNTC macrophages and observed that there is an increase in short 3’UTR-APA and IPA mRNA isoforms (Figure 6A and Supplementary Data 3). Box plot representations of 3’UTR-APA and IPA confirm the Volcano plots results and show that there is higher dispersion of values in IPA than in 3’UTR-APA (Figure 6B). Interestingly, *MBNL1*, a splicing regulator^78^ that was also shown to modulate alternative polyadenylation^79^, show a decrease in the distal PAS usage concomitant with an increase in proximal PAS use in the 3’UTR (Figure 6C). *CDC42*, which regulates macrophage chemotaxis and acts as an effector of phagocytosis^80, 81^, shows upregulation of an IPA mRNA isoform after *SRSF12* knockdown in macrophages (Figure 6D).

**Figure 6.**
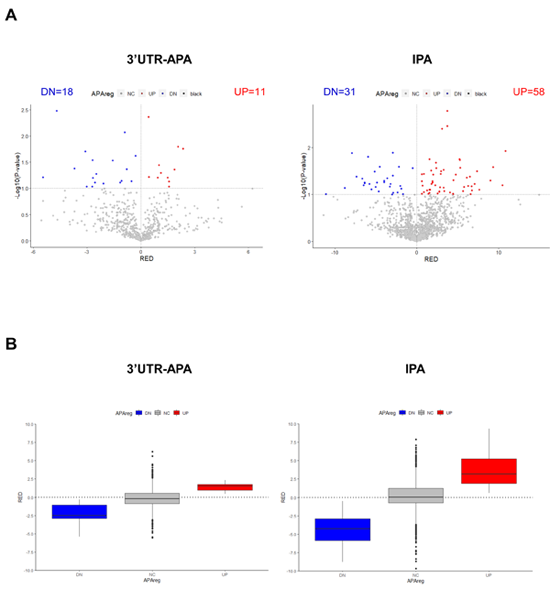
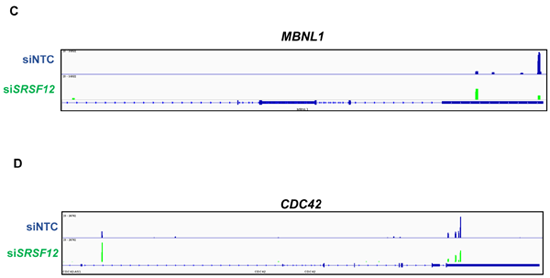
*SRSF12* knockdown induces 3’UTR-APA and IPA changes in M1 macrophages. **A.** Volcano plots of differential 3’UTR-APA (left) and IPA (right) in si*SRSF12 vs* siNTC in M1 macrophages. Red dots represent genes with RED >0, corresponding to upregulation of the lengthened isoform (A) or upregulated IPA(B). Blue dots represent genes with RED<0,, corresponding to upregulation of the shortened isoform (A) or downregulation of IPA (B), and p < 0.1, grey dots represent non-significantly expressed genes. **B.** Left, box plot with relative expression differences for 3’UTR-APA shortening (DN), 3’UTR lengthening (UP) and non-significant (NC) and right, Upregulated IPA (UP), Downregulated IPA (DN) and non-significant IPA (NC). The APAreg was calculated using a p-value<0.1 based on an unpaired t-test and a RED difference of 5%. **C.** *MBLN1* IGV gene profile and transcript counts in si*SRSF12* (**green**) and siNTC (**blue**) M1 macrophages shows upregulation of the proximal isoform upon SRSF12 knockdown. **D.** *CDC42* IGV gene profile and transcript counts in si*SRSF12* (**green**) and siNTC (**blue**) M1 macrophages shows upregulation of the IPA isoform upon SRSF12 knockdown.

## Discussion

Elucidating the gene expression and transcriptomic profile of pro-inflammatory macrophages, key immune cells at the tumor microenvironment, provides new insight into the mechanisms of immune evasion and cancer progression. However, the use of primary human macrophages in transcriptomic studies has been hampered by their highly plastic nature and thus the different methodologies for monocyte differentiation and macrophage polarization often result in diverse phenotypic outcomes. The use of monocytic cell lines, on the other hand, has the pitfall of not reproducing what occurs in primary cells. We used a robust and previously validated protocol for differentiation and polarization of primary human monocytes into a pro-inflammatory state and performed indirect co-cultures with two different CRC cell lines. We then performed an unbiased transcriptomic study where we assessed simultaneously gene expression, 3’UTR-APA and IPA events. Our 3’ RNA-Seq data report the biggest known catalogue of differential usage of alternative 3’ UTRs in primary human macrophages after M1 polarization, CRC stimuli and *SRSF12* knockdown, consisting of 30584 total 3’UTR-APA and IPA events, of which 1161 show significant differences. This data portfolio includes 3’UTR-APA and IPA events in key inflammatory-related genes, increasing the spectrum of known alterations in immune cells after pro-inflammatory polarization and exposure to cancer cells. M1 polarization induces substantial changes in macrophage gene expression (3005 genes out of a total of 19328 detected mRNAs, ie, 15.5%), which triples the 5% of DEGs previously reported in this population^39^. In agreement with a pro-inflammatory polarization, upregulated genes are involved in pro-inflammatory functions, while downregulated genes are related to anti-inflammatory and homeostatic functions.

Although pro-inflammatory macrophages are the main population in the colonic microenvironment at the beginning of the tumorigenic process and before immune evasion^8–10^, it has been suggested that during disease progression, macrophage profile may change towards a more anti-inflammatory and permissive microenvironment^82^. Our results are in agreement with these observations, as we show that after 24 hours of CRC co-culture macrophages present an upregulation of several anti-inflammatory genes. Single-cell analyses have identified TAM signatures in cancer^83^. After exposure to CRC cells for 24 hours, we identified our macrophage expression profile as belonging to the inflammatory cytokine-enriched TAMs (*G0S2*, *IL1B*, *IL6*, *S100A8*) and pro-angiogenic TAMs (*HES1*, *IL1B*, *IL8*, *S100A8*, *SERPINB2*, *THBS1*)

We identified a 35 DEG signature in co-cultured macrophages with upregulation of anti-inflammatory and pro-tumoral genes. Importantly, we observed that these results are in agreement with *in vivo* data from CRC patients, such as *LRP5*, *CXCL8* and *EREG*. In particular, *LRP5*, an member of the Wnt pathway^49^, which presents the highest increase in its expression in macrophages upon CRC co-culture, is overexpressed in tumor-associated macrophages (TAMs) in CRC patients. Moreover, an increase in *LRP5* expression correlates with patient survival as revealed by the Kaplan-Meier plot. These results provide new insight on a function for *LRP5* in the macrophage response to CRC.

Our 3’UTR-APA analyses reveal that M1 polarisation induces a strong 3’UTR-APA shortening and that affected genes have important functions in macrophage response to CRC, including in the Wnt and TGFβ pathways. We identified subsets of physiologically relevant genes that undergo differential 3’UTR-APA, including *IL17RA*, the receptor of the pro-inflammatory cytokine *IL17A* that plays important functions in inflammation and cancer^84^, and *TP53RK*, also known as *PRPK*, which codes for p53 protein kinase^85^. M1 polarization leads to a decrease in the usage of the most distal PAS and to an increase in the use of the most proximal PAS for *IL17RA* and *TP53RK*, which may affect their function. To our knowledge, this is the first study that identifies 3’UTR-APA mRNA isoforms after pro-inflammatory polarization of primary human macrophages.

Although changes in 3’UTR-APA have been described in CRC^86^ and in human macrophages infected with vesicular stomatitis virus^34^, our study describes for the first time how CRC exposure modifies the 3’UTR-APA profile of primary human macrophages. Co-culture with two distinct CRC cell lines, HCT-15 and RKO, allowed the identification of a subset of mRNA isoforms that are common to both cell lines. As these cell lines differ in several tumorigenic markers such as *APC*, *K*-*ras*, *B-raf*, *TGFBR2* and MLH1^42^, the identification of these new mRNA isoforms in response to CRC cells may provide insight on the crosstalk and signalling occurring between CRC cells and macrophages.

IPA is defined by the usage of an alternative PAS within an intron, as illustrated by the classical example of the membrane-bound and secreted IgM in B cells^87^. It has been shown that IPA is a widespread event^28^, in particular in immune cells^29^, and that it inactives tumor suppressor genes in leukaemia^30^. Here, we found that polarization induces IPA changes in macrophages and that there is a negative correlation between differential gene expression and IPA. We identified IPA events that may generate truncated proteins with other functions, as MAP3K8 and WDR33. MAP3K8 is necessary for activation of the MAPK/ERK pathway in macrophages, being key for producing the pro-inflammatory cytokine TNF-alpha (TNF) during immune responses. *MAP3K8* expresses an IPA mRNA isoform at low levels in M0, which is increased by 2-fold upon macrophage M1 polarization. Strikingly, this IPA isoform is even more expressed when macrophages are co-cultured with CRC. As this *MAP3K8* mRNA isoform is annotated and a possible protein is predicted, it is likely that this IPA isoform has implications for macrophages response to the tumor microenvironment. Of note, MAP3K8 full-length protein levels do not change upon polarization, but the production of a shorter protein is increased. WDR33 binds directly to the AAUAAA and thus has a key function in the mechanism of pre-mRNA 3’ end and processing by defining the PAS^62^. We found that M1 polarization leads to an increase in the production of a shorter IPA mRNA isoform, which is further increased upon CRC co-culture. It remains to be investigated the function of these short *MAP3K8* and *WDR33* isoforms in macrophages.

The expression of IPA mRNA isoforms expression has been shown to be modulated by Pcf11^31^ and it also depends on the competition between splicing and polyadenylation^88, 89^ that is illustrated by the membrane-bound *vs* secreted IgM^87^ and by the CT/CGRP mRNA isoforms^90^, and also by the function of U1A in polyadenylation^91, 92^. The efficiency of the splicing reactions is dictated by SR proteins^93^, which are classically viewed as splicing regulators albeit playing multiple roles in the different steps of mRNA biosynthesis. Here we show that *SRSF12/SRp35* (serine/arginine-rich splicing factor 12), which was initially described as an antagonist of SR proteins^77^, is highly upregulated after M1 polarization of primary human macrophages, and that *SRSF12* knockdown leads to a strong downregulation in global gene expression. Additionally, *SRSF12* depletion leads to 3’UTR-APA and IPA changes in a number of genes. One such example is *MBNL1*, a key splicing regulator that also regulates alternative polyadenylation^84^ and that undergoes proximal PAS usage upon *SRSF12* knockdown, and *CDC42*, a gene that encodes a small GTPase that regulates macrophage chemotaxis and phagocytosis^80, 81^, which presents an increase in IPA.

Overall, our findings show that pro-inflammatory polarization and CRC co-culture induces 3’UTR-APA shortening and IPA mRNA isoforms with a function in macrophage response to the microenvironment that may elicit a prompt response against the cancer challenge.

## Experimental Procedures

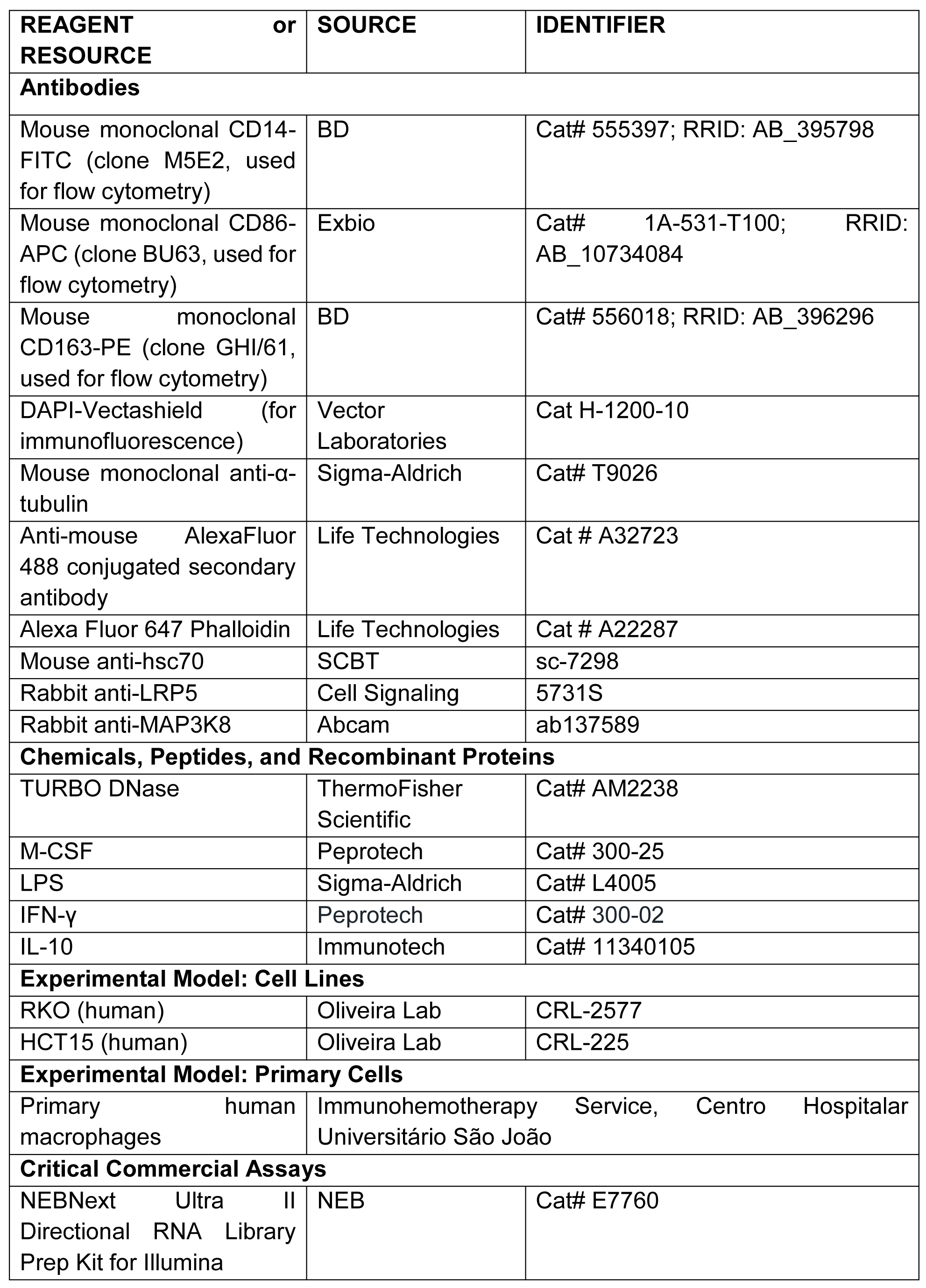
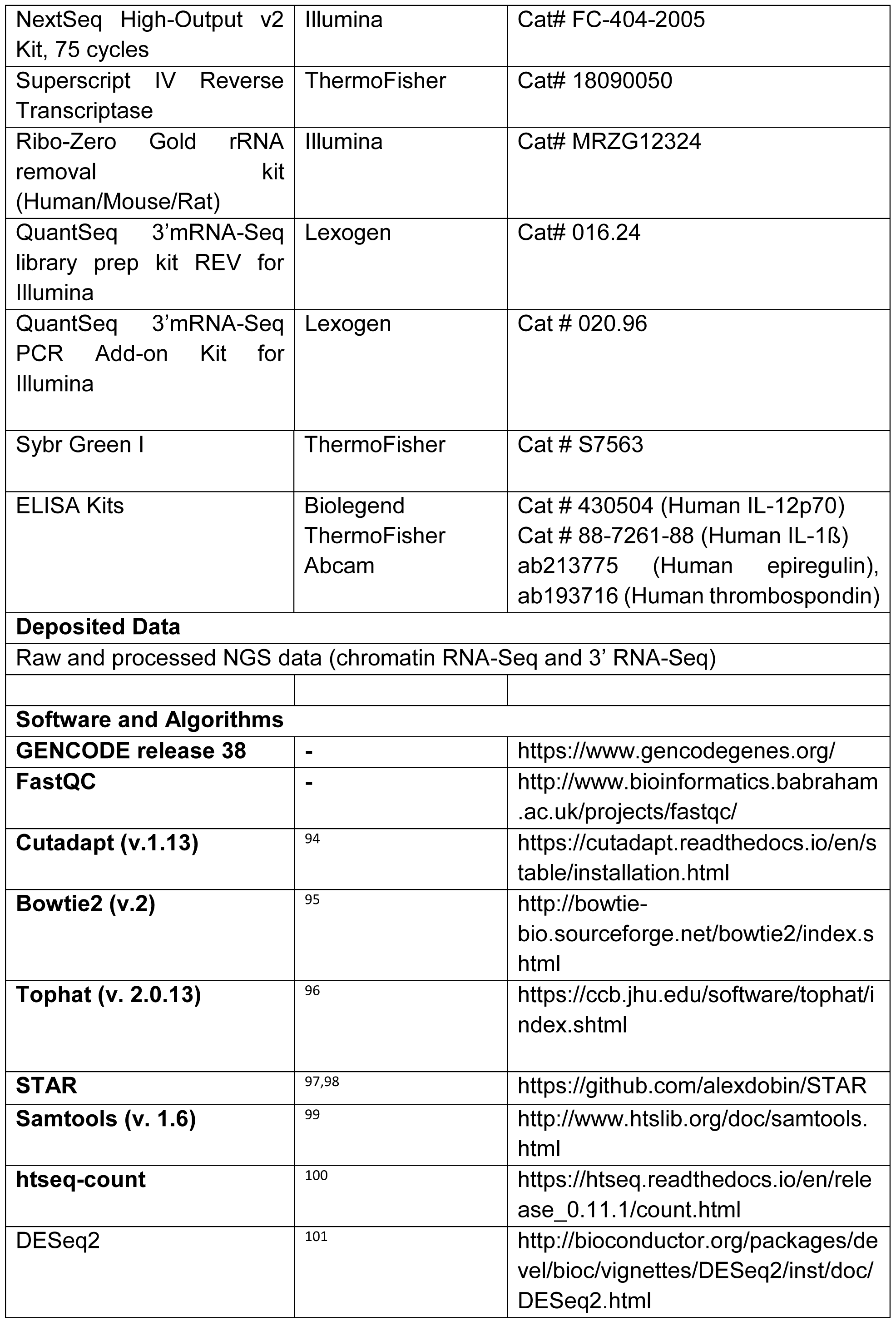
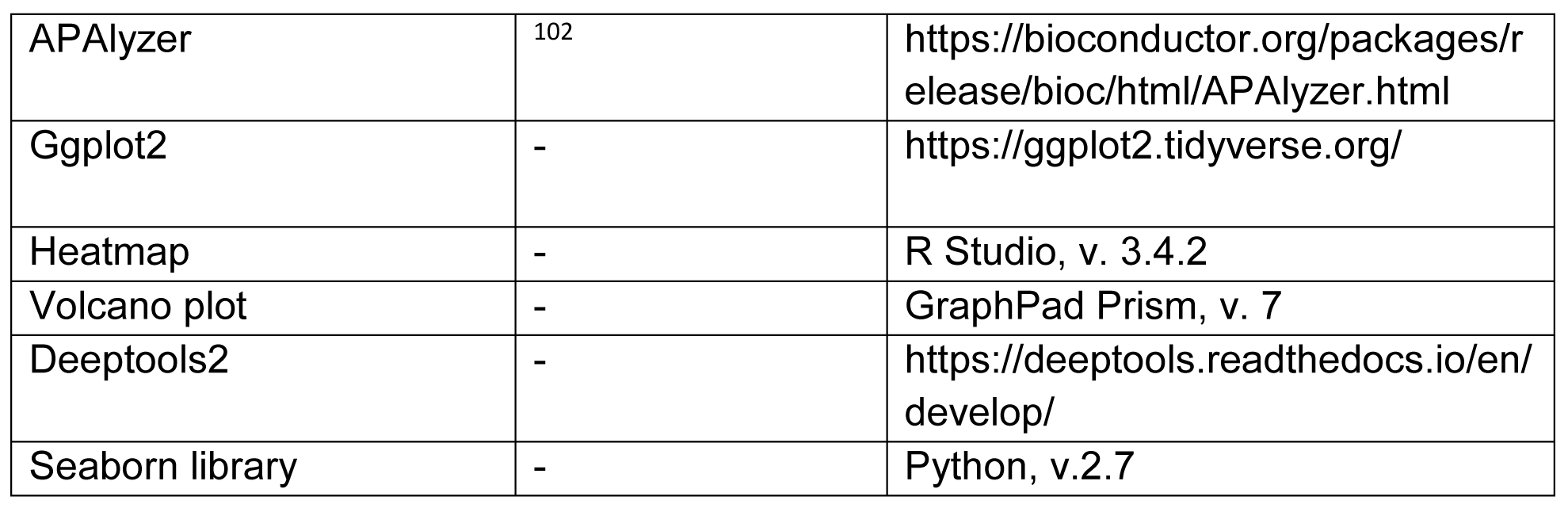

### Human primary macrophages

Human monocytes were isolated from anonymized buffy coats kindly provided by the Immunohemotherapy Service at Centro Hospitalar Universitário São João, Porto, Portugal. All experiments using human cells were conducted according to Portuguese and European guidelines for human research and approved by the institutional ethics committee (Protocol 90/19). Blood donors signed informed consent forms as part of the Ethics Committee guidelines. Healthy blood donors aged below 40 years old were selected for isolation.

## Methods Details

### Reagents

Macrophage colony stimulating factor (M-CSF, 300-25, Peprotech), lipopolysaccharide (LPS, L4005, Sigma-Aldrich), interferon-gamma (IFNγ, 300-02, Peprotech) and IL-10 (11340105, Immunotech) were prepared as in the manufacturers’ instructions. RPMI 1640 medium, streptomycin, penicillin, trypan blue and heat-inactivated fetal bovine serum (FBS) were obtained from Gibco. Bovine Serum Albumin (BSA) was obtained from nzytech.

### Cell lines

RKO and HCT-15 cells, both from American Type Culture Collection (ATCC) were maintained in complete Roswell Park Memorial Institute 1640 medium (RPMI) with GlutaMAX and supplemented with 10% FBS, 100 U/mL penicillin and 100 µg/mL streptomycin antibiotic solution. Cells were kept at 37°C, 5% CO2 in a humidified incubator, and subcultured every 3 to 5 days to maintain subconfluency. Trypan blue dye (Gibco) exclusion test was used to assess cell viability.

### Human monocyte isolation

Briefly, peripheral blood mononuclear cells (PBMCs) were isolated from buffy coats using Ficoll gradient centrifugation, followed by enrichment of CD14^+^ cells using CD14^+^ magnetic beads (Miltenyi Biotech, 130-050-201), according to the manufacturer instructions.

### Macrophage differentiation and activation

1×10^6^ human CD14^+^ cells were plated on 6-multiwell plates and maintained in RPMI 1640 medium supplemented with 10% FBS (Gibco), 100 U/mL penicillin (Invitrogen), 100 µg/mL streptomycin (Invitrogen) and 50 ng/mL M-CSF (Peprotech), and differentiated for 7 days. On day 7, media was refreshed and cells were polarized using 100 ng/mL LPS (Sigma-Aldrich) and 25 ng/mL IFNγ (Peprotech) in RPMI 1640 medium supplemented with 2% FBS without antibiotics for additional 24 h. Non-polarized M0 macrophages were maintained in RPMI 2% FBS without antibiotics. At the end of the experiment, spent culture media were collected for ELISA.

### Macrophage-cancer cell indirect co-cultures

24 hours after macrophage polarization, 3×10^5^ RKO or HCT-15 cells were seeded into transwell inserts with 1.0 µm pore size membrane (Corning 353102) and put on top of 1×10^6^ M1 macrophages. Co-cultures were maintained in complete RPMI 1640 medium for 24 h at 37° C and 5% CO2 in a humidified atmosphere, after which both cell populations and cell culture media were collected for downstream assays. M1 macrophages not exposed to co-culture were maintained in complete RPMI for the duration of the experiment.

### Macrophage siRNA knockdown

Briefly, 50 nM of a pool of 4 *SRSF12* siRNAs (Dharmacon ONTARGETplus SMARTpool siSRSF12) or 50 nM of a pool of a non-targeting siRNA control (Dharmacon ONTARGETplus SMARTpool siNTC) were transfected into 1.5×10^6^ M1 macrophages in 6-multiwells using 4 µL GenMute for Primary Human Macrophages (SignaGen) as a transfection agent, according to manufacturer’s instructions. Knockdown levels were assessed by RT-qPCR.

### Immunofluorescence

Macrophages grown on coverslips were fixed using 4% paraformaldehyde in 1X PBS for 15 min. Cells were quenched in 50 mM ammonium chloride for 10 min, permeabilized in 0.2 % Triton X-100 in 1x PBS for 5 min and incubated in 5% BSA blocking solution for 30 min, all at room temperature, before incubating for 1 h with mouse anti-ß-tubulin and for 45 min with anti-mouse AlexaFluor 488 secondary antibody, protecting from light. Coverslips were then incubated with Phalloidin-FITC for 15 min at room temperature, and mounted with DAPI-Vectashield. Confocal images were acquired in a Leica Spectral Confocal TCS-SP5 AOBS (Wetzlar, Germany) microscope, using the 40x or 63x oil objectives. All experimental conditions were surveyed using the same microscope conditions. Images were analyzed and processed using the FIJI package for Image J^103^.

### Flow cytometry

Macrophages were incubated with Accutase (eBioscience, Asymetrix) for 30 min at 37 °C, harvested by gentle scraping, washed with 1x PBS and resuspended in FACS buffer (1x PBS, 2% FBS, 0.02% sodium azide), blocked with FACS blocking reagent and resuspended in diluted antibodies for 30 min at 4°C in the dark: CD14-FITC (BD 555397, clone M5E2, 1:50 dilution), CD86-APC (Exbio 1A-531, clone BU63, 1:25 dilution), and CD163-PE (BD 556018, clone GHI/61, 1:25 dilution). Unstained cells were used as control. Cell surface markers were detected on FACS Canto II (BD Biosciences), using FACS Diva® software, and data were analyzed using FlowJo software v.10.4.

### in silico analyses

Nucleotide sequences and mRNA profiles were obtained from NCBI and Ensembl genome browsers. The Ensembl and UCSC genome browsers and the APAdb and Aceview databases were accessed to search for alternative polyadenylation sites on the 3’ UTR of human genes.

### RNA extraction

Cells were washed with ice-cold PBS and resuspended in 500 µL TRIzol (Invitrogen) to extract total RNA according to manufacturer’s instructions, with the exception that all samples were incubated overnight at −80°C during isopropanol precipitation. Total RNA quantity was determined with a Nanodrop 1000 spectrometer (Thermo Fisher Scientific) or a Qubit Fluorometer, using Qubit RNA HS Assay Kit (ThermoFisher).

### cDNA synthesis and Quantitative Real-Time PCR (RT-qPCR)

Total RNA was digested with DNase I (Roche) for 25 min at 37°C, inactivating the enzyme for 10 min at 80°C. RNA was reverse transcribed using SuperScript IV (ThermoFisher) and random hexamers, according to manufacturer’s instructions. RT-qPCR reactions were performed in triplicate using SYBR Select Master Mix (Applied Biosystems) following the manufacturer’s protocol and 0.125 µM of the primer pairs listed in Supplementary Table 1. Negative controls without reverse transcriptase were used to detect possible gDNA contamination. Relative expression was calculated relative to the reference gene *18S*. For quantification of the relative expression of *CCR7* and *TGFB*, RT-qPCR reactions were performed in triplicates using TaqMan Probes and Master Mix (Applied Biosystems). Relative expression was calculated relative to the reference gene *ACTB*. Results were analyzed applying the ΔΔCt method^104^.

### Primer Design

For RT-qPCR of APA candidates, primers were designed to the coding and most distal PAS (pA2), as previously described^105, 106^. Primer sequences used in the RT-qPCRs are listed on Supplementary Table 1.

### Chromatin-bound fractionation

Total RNA from macrophages was fractionated as described in^38^. Briefly, 6×10^6^ cells for each condition were washed twice with ice-cold 1x PBS, spun down, and cell pellets were lysed with HLB/NP40 buffer (10 mM Tris-HCl pH 7.5 / 10 mM NaCl / 2.5 mM MgCl2 / 0.05% NP-40 Igepal (Sigma)), carefully underlaid with HLB/NP40/Sucrose (HLB/NP-40 + 25% sucrose) and centrifuged at 208 *g* for 5 minutes at 4°C to isolate the nuclei from the cytoplasm. The chromatin fraction was isolated via treatment with NUN1 solution (20 mM Tris-HCl pH 7.9 / 75 mM EDTA/ 50% v/v glycerol / Protease Inhibitor cOmplete 1x (Roche) added fresh) and NUN2 buffer (20 mM HEPES-KOH pH 7.6 / 300 mM NaCl / 0.2 mM EDTA / 7.5 mM MgCl2 / 1% v/v Igepal / 1 M Urea), with 15 min incubating on ice, followed by centrifugation at 15680 *g* for 10 minutes at 4°C, separating from the nucleoplasm supernatant. The RNA-bound chromatin pellet was then resuspended in high salt buffer HSB (10 mM Tris-HCl pH 7.5 / 500 mM NaCl / 10 mM MgCl2) and digested with TURBO DNase (Invitrogen) and Proteinase K (10 mg/mL; Ambion) for 10 min each at 37°C and 182 *g* for 10 min. All buffers were ice-cold. RNA was then extracted from the cytoplasmic and chromatin-bound fractions with Trizol. To ensure adequate fraction separation was achieved, *MALAT1* RNA levels were measured by RT-qPCR in cytoplasmic and chromatin-bound fractions; *MALAT1* presence was only detected in the chromatin-bound fraction.

## RNA-Seq

### Library preparation

For 3’ mRNASeq (3’ Seq), polyA^+^ mRNA was obtained from total RNA using Dynabeads mRNA purification kit (ThermoFisher), according to manufacturer’s protocol. QuantSeq Rev (Lexogen) mRNA libraries were generated at i3S, Porto, Portugal, using the manufacturer’s protocol, quantified using Agilent TapeStation and KAPA Library Quantification Kit (KAPA Biosystems), and sequenced with NextSeq500 using single-end 75-nucleotide reads in the Lexogen Sequencing Facility (Vienna). 3’RNA-Seq of si*SRSF12* and siNTC was performed with total RNA.

For ChrRNA-Seq, rRNAs were removed using Ribo-Zero Gold rRNA removal Kit (Human/Mouse/Rat) (Illumina) according to manufacturer’s protocol. Chromatin-bound RNA libraries from three biological replicates were generated using NEBNext Ultra II Directional Prep Kit for Illumina (New England Biolabs), according to the manufacturer’s protocol, and sequenced with Illumina NextSeq High-Output Kit, 75 cycles, using paired-end 42-nucleotide reads.

#### Data pre-processing

In all datasets, quality control was performed using FastQC and adapters were trimmed with Cutadapt (v1.13), discarding reads under 10 bases.

For 3’RNA-Seq, raw sequencing data were filtered to detect accurate poly(A) signals, since this sequencing technique may have intrinsic leakiness in polyadenylation signal discovery. Briefly, the input fastq file was mapped to the hg38 reference and three filters were applied in sequence: 1) a Transcript Termination Site (TTS) filter, removing transcripts not lying within ∼10 nt of an annotated transcript end site; 2) a Motif filter removed all sequencing peaks whose read start contained a subset of hexamer sequences upstream^107^ and 3) a downstream A content filter checked the amount of As in a window downstream of the read start; if a given peak passes the validated threshold, the read is considered internal priming and is removed. All reads passing the three filters are retained as non-internal priming. Total fragments passing all filters ranged from 47.4-70.7% (Supplementary Table 2. Data were aligned to the reference human genome (hg38) using STAR^108^. Aligned data had 75.6%-85.4% overall read mapping rate (Supplementary Table 2).

For ChrRNA-Seq, data were aligned to the reference human genome (hg38) using TopHat (v.2.1)^96^, allowing read pairs to be separated by 3 kb and only one alignment to the reference genome for each read. Aligned data had 89%-91.2% overall read mapping rate (Supplementary Table 3).

In all datasets, aligned reads were processed with SAMtools^98^ and counted with htseq-count^100^ and gene expression differences were calculated using DESeq2^101^. Sequencing read distribution through the genome was visualized in IGV, v 7.0, using BedGraphToBigWig.

To increase statistical robustness to our analysis, we only considered genes with >5 mean TPM, Log2Fold Changes ≥ 1 or ≤ -1, and adjusted p-value < 0,05, calculated by DESeq2^109^. Additionally, to establish data set reproducibility, we used, used 3 independent experiments with macrophages from three donors and analyzed the Pearson correlation values. R values exceeded 0.9 for each condition, exceeded 0.8 between pro-inflammatory and co-cultured macrophages, and exceeded 0.5 between M0 and M1 (Figure S1G).

mRNA isoform expression differences were calculated in the 3’RNA-Seq data using APAlyzer to generate lists of genes showing differential expression of alternative mRNA isoforms by alternative polyadenylation. APAlyzer is a Bioconductor package which analyses 3’UTR-APA and IPA based on annotated, conserved polyA sites at the recommended cutoff of relative expression between conditions (5%), using p-value<0.05 with the exception of siSRSF12 3’RNA-Seq data where p<0.1. For downstream analyses, we focused on isoforms of protein-coding genes as per ENSEMBL databases, excluding out-of-scope isoforms such as NMD.

To clarify the identity of each mRNA isoform being produced in each condition, mRNA isoforms detected by 3’ RNA-Seq were compared to Chr-bound RNA-Seq data from M1 and M1 + RKO macrophages and to transcripts deposited in the ENSEMBL database.

For SRSF12 knockdown 3’RNA-Seq, the bioinformatic pipeline was the same, with the exception that the initial filtering algorithm was not applied.

#### Candidate gene selection

Criteria for 3’RNA-Seq candidate gene validation were as follows: DGE candidates were selected for large fold change differences between control and co-cultured macrophages, robustness of response after macrophage co-culture with both CRC cell lines, and relevance to macrophage biology. mRNA isoform candidates were selected according to the following criteria: i) prevalent 3’ UTR mRNA isoform shift between the 3’ RNA-Seq signal in the two mRNA isoforms with the highest reads, ii) robustness of response after macrophage polarization and co-culture with both cell lines (increase in proximal or distal mRNA isoform) and iii) relevance to macrophage biology and inflammation.

#### TCGA analysis of patient survival

Datasets for colon carcinoma and normal tissue with CD68^high^ expression were retrieved from the Cancer Genome Atlas (TCGA^110^) and analyzed for post-diagnosis survival data, using TIMER. The normal tissue median values were used as threshold to define “high” and “low” expression levels for each identified DEGs in cancer patients, quantified as FPKM (fragments per Kilobase million). The correlation index (R) indicated a moderate correlation if between 0.25 and 0.35, and a strong correlation if higher than 0.35.

#### Gene Ontology (GO) and Gene Set Enrichment Analysis (GSEA)

Gene Ontology was performed using PANTHER v. 16 (http://www.pantherdb.org/). Functional annotation enrichment was performed on the genes with differential expression using GSEA software (Broad Institute).

#### Enzyme-Linked Immunosorbent Assay (ELISA)

Cytokines and metabolites were quantified in the culture media of mono and co-cultures by ELISA using kits from ThermoFisher and BioLegend, according to manufacturer’s instructions.

#### Western Blot

Macrophage lysates were prepared with RIPA buffer (50 mM Tris-HCl pH 7.5 / 1% NonidetP (NP)-40 / 150 mM NaCl / 2 mM EDTA) with protease inhibitors (Protease Inhibitor cOmplete 1x). Protein concentration was determined through the Bradford method and 25 µg of protein was loaded and run in a 7.5% (for LRP5) or 10% acrylamide gel, which was subsequently transferred into nitrocellulose membranes (GE Healthcare), blocked with 5% non-fat powdered milk or BSA in TBS + 0.1 % Tween 20 (TBS-T) for 1 hour, and incubated overnight with primary antibodies at 4°C (Methods Table). HRP-conjugated secondary antibodies (SCBT) were incubated for 1 hour at RT. Signal was detected by incubation with ECL Substrate (GE Healthcare), using the Chemidoc Imaging System (BioRad) and band intensity was measured in the ImageLab Software (BioRad).

#### Statistical analysis

All presented data is mean ± standard deviation (SD). Data was evaluated for normal distribution through the Kolmogorov-Smirnov test. Comparisons between two independent groups were performed using the paired Student’s t-test or the Wilcoxon test. Except where otherwise stated, p-values <0.05 were considered statistically significant (* = p < 0.05; ** = p < 0.01; *** = p < 0.005; **** = p < 0.0001). Graphs were made in the GraphPad Prism software, using version 7.0a for Mac, GraphPad Software, La Jolla California USA, www.graphpad.com.

## Acknowledgements

We thank members of the Gene Regulation and Tumor and Microenvironment Interactions groups at i3S, and the Proudfoot’s and Murphy’s laboratories at the University of Oxford, for critical discussions. The authors acknowledge the Immunohemotherapy Service at Centro Hospitalar Universitário São João for their kind donation of the buffy coats needed for this study. The authors also acknowledge the i3S Scientifics Platforms: María Lázaro from the Bioimaging unit (member of the Portuguese Platform of Bioimaging - PPBI-POCI-01-0145-FEDER-022122), Paula Magalhães and Tânia Meireles from the Cell Culture and Genotyping unit, Mafalda Rocha and Rob Mensink from the Genomics Unit, and Catarina Meireles from the Translational Cytometry Unit. JW also acknowledges Bruno Pereira for help with the GSEA software (Broad Institute).

## Funding

This work was supported by the “Cancer Research on Therapy Resistance: From Basic Mechanisms to Novel Targets”—NORTE-01–0145-FEDER-000051 project, supported by Norte Portugal Regional Operational Programme (NORTE 2020), under the PORTUGAL 2020 Partnership Agreement, through the European Regional Development Fund (ERDF), by the European Union’s Horizon 2020 research and innovation programme under grant agreement No 952334, by FCT under the project EXPL/SAU-PUB/1073/2021 and by Programa Operacional Regional do Norte and co-funded by European Regional Development Fund under the project “The Porto Comprehensive Cancer Center” with the reference NORTE-01–0145-FEDER-072678—Consórcio P.CCC—Porto Comprehensive Cancer Center to AM, to NJP (Wellcome Trust Investigator Award [107928/Z/15/Z] and ERC Advanced [339270] grants), to IPC (DL 57/2016/CP1355/CT0016) and to JW by FCT/GABBA PhD fellowship (PD/BD/114168/2016).

## Ethical Statement

Buffy coats are waste-products from healthy blood donations that contain high amounts of leukocytes, which are isolated in accordance with protocol 90/19 approved by the Centro Hospitalar Universitário São João Ethics Committee. Anonymized donors filled informed consent forms prior to each blood donation.

## Supplementary figures

**Supplementary Figure S1.**
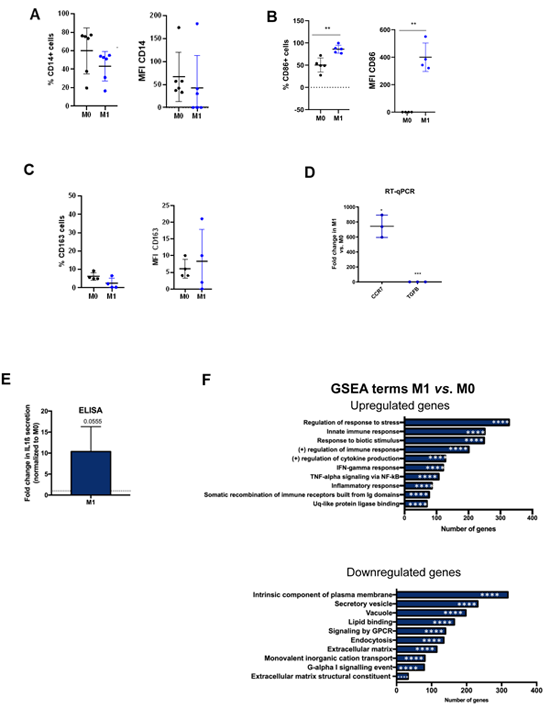
**A-E**. Inflammatory profile characterization of M1 normalized to M0. **A-C**. Percentage of positive cells for CD14 (**A**, n=6), CD86 (**B**, n=6) and CD163 (**C**, n=4) **D**. Pro-inflammatory marker *CCR7* and anti-inflammatory marker *TGFB* measured by RT-qPCR (n=3). **E**. IL1ß secretion in M1 *vs*. M0 macrophages, measured by ELISA (n=3). A-D Student’s t-test: * = p < 0,05; ** = p < 0,01. M0 in black, M1 in blue, M2 in light grey**. F**. GSEA terms of genes upregulated (top) and downregulated (bottom) in M1 *vs*. M0. **** = p < 0,0001.

**Supplementary Figure S2.**
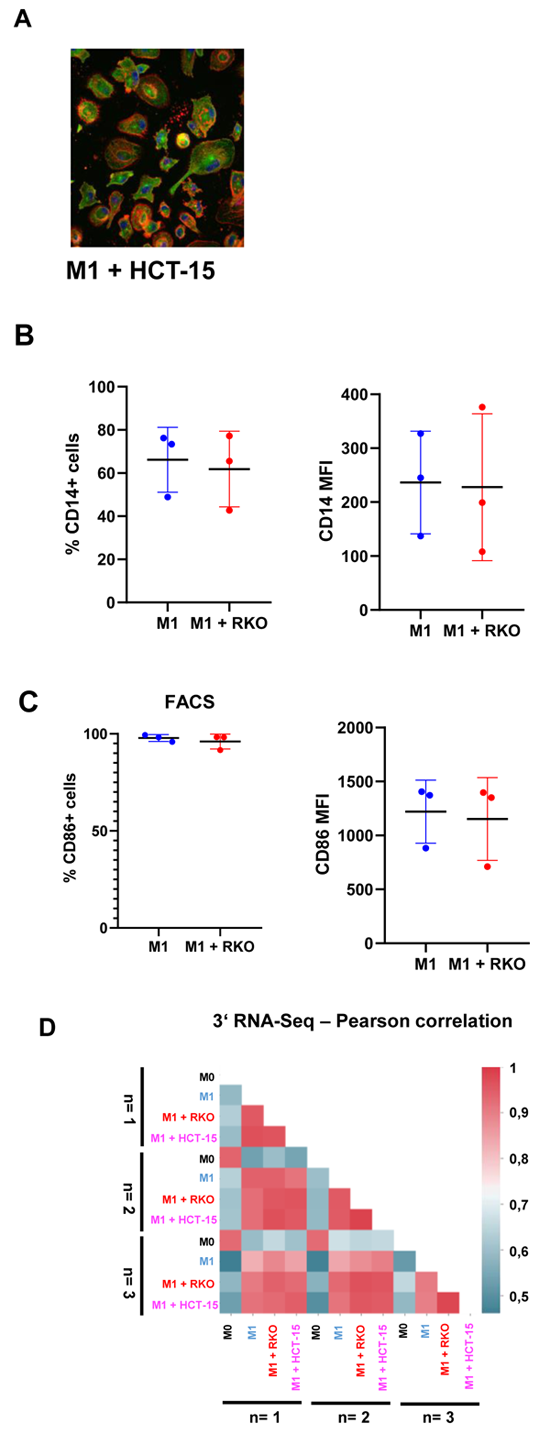
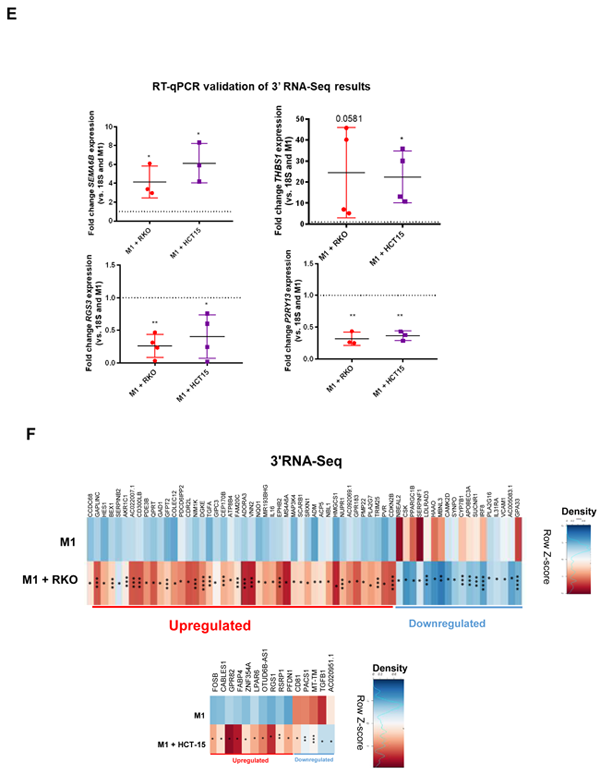
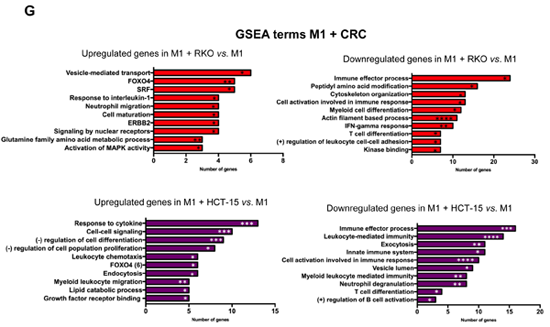
**A.** Confocal image of macrophages co-cultured with CRC cell line HCT-15 for 24 hours as shown in Figure 1B (M1 + HCT-15). DNA stained in blue, actin stained in red, tubulin stained in green. **B-C**. Inflammatory profile characterization of M1 + RKO *vs*. M1. Percentage of positive cells (left) and median fluorescence intensity (right) for CD14 (B), and CD86 (C) measured by flow cytometry in M1+RKO (**red**) *vs*. M1 (**blue**). **D**. Heatmap of Pearson correlation coefficients for 3’ mRNA-Seq samples, using transcript counts per million (CPM); Scale: 0-1, 0 = no correlation between populations; 1= total correlation between populations (n=3). **E**. RT-qPCR validation of common 3‘RNA-Seq DEGs in co-cultured macrophages. Top: upregulated genes *SEMA6B* (n=3), *THBS1* (n=4), Bottom: downregulated genes *RGS3* (n=4) and *P2RY13* (n=3). p-value calculated using Student’s t-test * = p < 0,05; ** = p < 0,01. **F.** Representative donor heatmap of genes exclusively upregulated in M1 + RKO *vs*. M1 (left) M1 + HCT-15 *vs.* M1 (right), ordered by fold change (n=3, p-value calculated using DESeq2 test through the Benjamini-Hochberg method. * = p < 0,05; ** = p < 0,01; *** = p < 0,005; **** = p<0,0001). **G**. Top 10 GSEA terms of upregulated genes (left) and downregulated genes (right) in M1 + RKO *vs.* M1 (left) and M1 + HCT-15 *vs.* M1 (right). n= 3, p-value calculated using GSEA: * = p < 0,05; ** = p < 0,01; *** = p < 0,005; **** = p<0,0001.

**Supplementary Figure S3.**
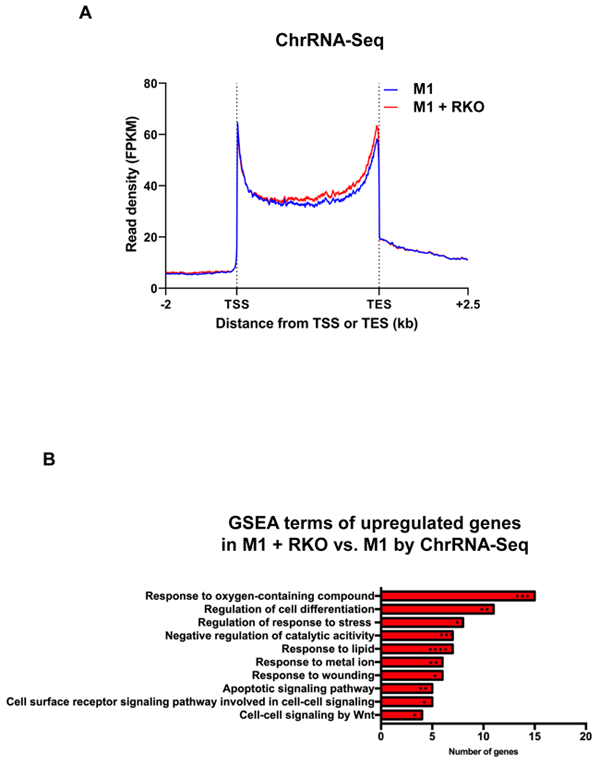
**A.** Metaprofile of protein-coding genes of ChrRNA-Seq data, from -2kb upstream of the transcription start site (TSS) to 2.5 kb downstream of the transcription end site (TES). **B**. GSEA terms of genes upregulated in ChrRNA-Seq data. p-value calculated using GSEA: * = p < 0,05; ** = p < 0,01; *** = p < 0,005; **** = p<0,0001.

**Supplementary Table S1.**
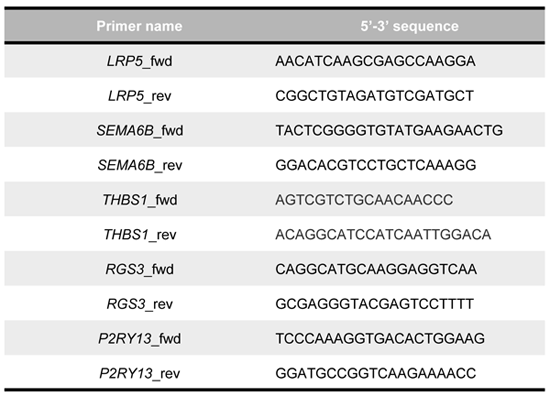
Primer pairs used for RT-qPCR validation.

**Supplementary Table S2.**
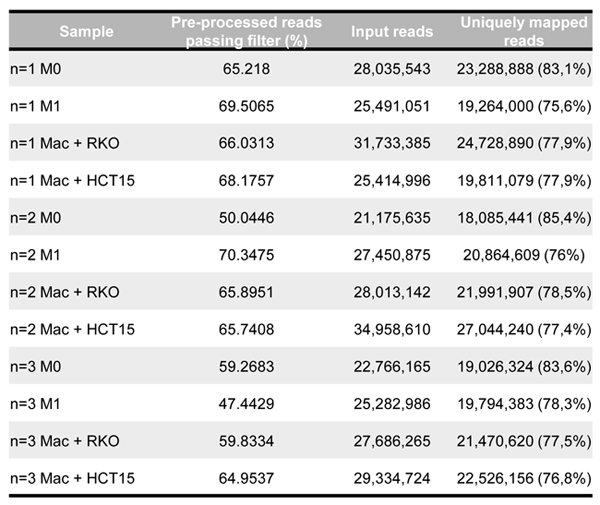
Percentage of reads passing filters after pre-processing, respective input read and uniquely mapped reads in 3’ RNA-Seq of M0, M1 and co-cultured macrophages data.

| Sample | Pre-processed reads passing filter (%) | Input reads | Uniquely mapped reads |
| --- | --- | --- | --- |
| n=1 M0 | 65.218 | 28,035,543 | 23,288,888 (83,1%) |
| n=1 M1 | 69.5065 | 25,491,051 | 19,264,000 (75,6%) |
| n=1 Mac + RKO | 66.0313 | 31,733,385 | 24,728,890 (77,9%) |
| n=1 Mac + HCT15 | 68.1757 | 25,414,996 | 19,811,079 (77,9%) |
| n=2 M0 | 50.0446 | 21,175,635 | 18,085,441 (85,4%) |
| n=2 M1 | 70.3475 | 27,450,875 | 20,864,609 (76%) |
| n=2 Mac + RKO | 65.8951 | 28,013,142 | 21,991,907 (78,5%) |
| n=2 Mac + HCT15 | 65.7408 | 34,958,610 | 27,044,240 (77,4%) |
| n=3 M0 | 59.2683 | 22,766,165 | 19,026,324 (83,6%) |
| n=3 M1 | 47.4429 | 25,282,986 | 19,794,383 (78,3%) |
| n=3 Mac + RKO | 59.8334 | 27,686,265 | 21,470,620 (77,5%) |
| n=3 Mac + HCT15 | 64.9537 | 29,334,724 | 22,526,156 (76,8%) |

**Supplementary Table S3.**
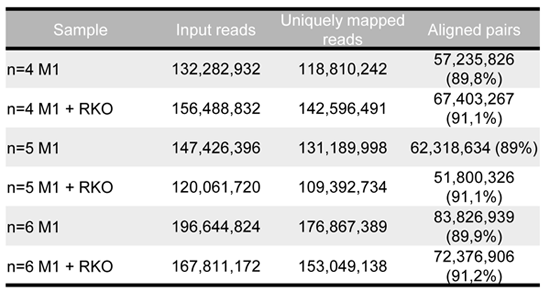
Percentage of reads passing filters after pre-processing, respective input read and uniquely mapped reads in ChrRNA-Seq data.

**Supplementary Table S1** – Percentage of reads passing filters after pre-processing, respective input and uniquely mapped reads in 3’ RNA-Seq of M0, M1 and co-cultured macrophages data.

**Supplementary Table S2** – Primer pairs used for RT-qPCR validation.

**Supplementary Table S3** – Percentage of reads passing filters after pre-processing, respective input and uniquely mapped reads in ChrRNA-Seq of M1 and co-cultured macrophages data.

**Supplementary Data 1** – Fold change and p-value of genes with differential expression calculated by DESeq2 in M1 *vs*. M0 macrophages, M1+RKO *vs*. M1 and M+HCT15 *vs*. M1.

**Supplementary Data 2** – Relative expression differences and p-value of genes with APA and IPA differences calculated by Apalyzer in M1 *vs*. M0 macrophages, M1+RKO *vs*. M1 and M+HCT15 *vs*. M1.

**Supplementary Data 3** – Fold change and p-value of genes with differential expression calculated by DESeq2, 3’UTR-APA and IPA calculated by Apalyzer, in M1 si*SRSF12 vs*. M1 siNTC macrophages.

## Notes

### Competing Interest Statement

The authors have declared no competing interest.

## References

1. WHO. Estimated number of cases worldwide, both sexes, all ages. 160 (2018).

2. Pinto, M. L. et al. The Two Faces of Tumor-Associated Macrophages and Their Clinical Significance in Colorectal Cancer. Front Immunol 10, 1–12 (2019).

3. Mantovani, A. & Locati, M. Tumor-associated macrophages as a paradigm of macrophage plasticity, diversity, and polarization lessons and open questions. Arterioscler Thromb Vasc Biol 33, 1478–1483 (2013).

4. Locati, M., Mantovani, A. & Sica, A. Macrophage Activation and Polarization as an Adaptive Component of Innate Immunity. Advances in Immunology vol. 120 (2013).

5. Hanahan, D. & Weinberg, R. A. The Hallmarks of Cancer. 100, 57–70 (2000).

6. Hanahan, D. & Weinberg, R. A. Hallmarks of cancer: The next generation. Cell 144, 646–674 (2011).

7. Németh, Z. H. et al. Adenosine Augments IL-10 Production by Macrophages through an A 2B Receptor-Mediated Posttranscriptional Mechanism . The Journal of Immunology 175, 8260– 8270 (2005).

8. Mantovani, A., Sica, A., Allavena, P., Garlanda, C. & Locati, M. Tumor-associated macrophages and the related myeloid-derived suppressor cells as a paradigm of the diversity of macrophage activation. Hum Immunol 70, 325–330 (2009).

9. He, Z. & Zhang, S. Tumor-Associated Macrophages and Their Functional Transformation in the Hypoxic Tumor Microenvironment. Front Immunol 12, 1–11 (2021).

10. Yahaya, M. A. F., Lila, M. A. M., Ismail, S., Zainol, M. & Afizan, N. A. R. N. M. Tumour-Associated Macrophages (TAMs) in colon cancer and how to reeducate them. J Immunol Res 2019, (2019).

11. Lawrence, T. & Natoli, G. Transcriptional regulation of macrophage polarization: Enabling diversity with identity. Nat Rev Immunol 11, 750–761 (2011).

12. Tian, B. & Manley, J. L. Alternative cleavage and polyadenylation: The long and short of it. Trends Biochem Sci 38, 312–320 (2013).

13. Braz, S. O. et al. Expression of Rac1 alternative 3′ UTRs is a cell specific mechanism with a function in dendrite outgrowth in cortical neurons. Biochim Biophys Acta Gene Regul Mech 1860, 685–694 (2017).

14. Berkovits, B. D. & Mayr, C. Alternative 3’ UTRs act as scaffolds to regulate membrane protein localization. Nature 522, 363–367 (2015).

15. Pai, A. A. et al. Widespread Shortening of 3’ Untranslated Regions and Increased Exon Inclusion Are Evolutionarily Conserved Features of Innate Immune Responses to Infection. PLoS Genet 12, 1–24 (2016).

16. Matsui, S. I., Weinfeld, H. & Sandberg, A. A. Quantitative conservation of chromatin-bound RNA polymerases I and II in mitosis. Implications for chromosome structure. Journal of Cell Biology 80, 451–464 (1979).

17. Mayr, C. Regulation by 3′–Untranslated Regions. Annu Rev Genet 51, annurev-genet-120116-024704 (2017).

18. Castello, A. et al. Insights into RNA Biology from an Atlas of Mammalian mRNA-Binding Proteins. Cell 149, 1393–1406 (2012).

19. Elkon, R., Ugalde, A. P. & Agami, R. Alternative cleavage and polyadenylation: extent, regulation and function. Nat Rev Genet 14, 496–506 (2013).

20. Tian, B. & Manley, J. L. Alternative polyadenylation of mRNA precursors. Nat Rev Mol Cell Biol 18, 18–30 (2016).

21. Derti, A. et al. A quantitative atlas of polyadenylation in five mammals. Genome Res 22, 1173–1183 (2012).

22. Shi, Y. Alternative polyadenylation: new insights from global analyses. RNA 18, 2105–17 (2012).

23. Tian, B., Hu, J., Zhang, H. & Lutz, C. S. A large-scale analysis of mRNA polyadenylation of human and mouse genes. Nucleic Acids Res 33, 201–212 (2005).

24. Wang, E. T. et al. Alternative isoform regulation in human tissue transcriptomes. Nature 456, 470–476 (2008).

25. Ma, W. & Mayr, C. A Membraneless Organelle Associated with the Endoplasmic Reticulum Enables 3′UTR-Mediated Protein-Protein Interactions. Cell 175, 1492–1506.e19 (2018).

26. Mitra, M., Johnson, E. L. & Coller, H. A. Alternative polyadenylation can regulate post-translational membrane localization. Trends Cell Mol Biol 10, 37–47 (2016).

27. Pereira-Castro, I. & Moreira, A. On the function and relevance of alternative 3′-UTRs in gene expression regulation. Wiley Interdiscip Rev RNA 1–30 (2021) doi:10.1002/wrna.1653.

28. Tian, B., Pan, Z. & Ju, Y. L. Widespread mRNA polyadenylation events in introns indicate dynamic interplay between polyadenylation and splicing. Genome Res 17, 156–165 (2007).

29. Singh, I. et al. Widespread intronic polyadenylation diversifies immune cell transcriptomes. Nat Commun 9, (2018).

30. Lee, S. H. et al. Widespread intronic polyadenylation inactivates tumour suppressor genes in leukaemia. Nature 561, 127–131 (2018).

31. Wang, R., Zheng, D., Wei, L., Ding, Q. & Tian, B. Regulation of Intronic Polyadenylation by PCF11 Impacts mRNA Expression of Long Genes. Cell Rep 26, 2766–2778.e6 (2019).

32. Sandberg, R., Neilson, J. R., Sarma, A., Sharp, P. a & Burge, C. B. Proliferating cells express mRNAs with shortened 3’ UTRs and fewer microRNA target sites. Science (1979) 320, 1643– 1647 (2008).

33. Mayr, C. & Bartel, D. P. Widespread Shortening of 3’UTRs by Alternative Cleavage and Polyadenylation Activates Oncogenes in Cancer Cells. Cell 138, 673–684 (2009).

34. Jia, X. et al. The role of alternative polyadenylation in the antiviral innate immune response. Nat Commun 8, 14605 (2017).

35. Nogueira, E. et al. Neutral PEGylated liposomal formulation for efficient folate-mediated delivery of MCL1 siRNA to activated macrophages. Colloids Surf B Biointerfaces 155, 459–465 (2017).

36. Wilton, J. et al. Simultaneous studies of gene expression and alternative polyadenylation in primary human immune cells. Methods in Enzymology vol. mRNA 3’ En (Elsevier Inc., 2021).

37. Ohradanova-Repic, A., Machacek, C., Fischer, M. B. & Stockinger, H. Differentiation of human monocytes and derived subsets of macrophages and dendritic cells by the HLDA10 monoclonal antibody panel. Clin Transl Immunology 5, (2016).

38. Nojima, T., Gomes, T. T., Carmo-Fonseca, M. & Proudfoot, N. J. Mammalian NET-seq analysis defines nascent RNA profiles and associated RNA processing genome-wide. Nat Protoc 11, 413–428 (2016).

39. Martinez, F. O., Gordon, S., Locati, M. & Mantovani, A. Transcriptional profiling of the human monocyte-to-macrophage differentiation and polarization: new molecules and patterns of gene expression. Journal of Immunology 177, 7303–11 (2006).

40. Pinto, A. T. et al. Intricate macrophage-colorectal cancer cell communication in response to radiation. PLoS One 11, 1–20 (2016).

41. Larionova, I. et al. Tumor-Associated Macrophages in Human Breast, Colorectal, Lung, Ovarian and Prostate Cancers. Front Oncol 10, 1–34 (2020).

42. Ahmed, D. et al. Epigenetic and genetic features of 24 colon cancer cell lines. Oncogenesis 2, e71 (2013).

43. Haderk, F., et al. Tumor-derived exosomes modulate PD-L1 expression in monocytes. Sci Immunol 2, eaah5509 (2017).

44. Ruiz-López, L. et al. The role of exosomes on colorectal cancer: A review. J Gastroenterol Hepatol 33, 792–799 (2018).

45. Kaler, P., Augenlicht, L. & Klampfer, L. Macrophage-derived IL-1Β stimulates Wnt signaling and growth of colon cancer cells: A crosstalk interrupted by vitamin D“3. Oncogene 28, 3892– 3902 (2009).

46. Gelfo, V. et al. Roles of il-1 in cancer: From tumor progression to resistance to targeted therapies. Int J Mol Sci 21, 1–14 (2020).

47. Abdelwahab, E. M. M. et al. Wnt signaling regulates trans-differentiation of stem cell like type 2 alveolar epithelial cells to type 1 epithelial cells. Respir Res 20, 1–9 (2019).

48. Kim, M. et al. Understanding the functional role of genistein in the bone differentiation in mouse osteoblastic cell line MC3T3-E1 by RNA-seq analysis. Sci Rep 8, 1–12 (2018).

49. Logan, C. Y. & Nusse, R. The Wnt signaling pathway in development and disease. Annu Rev Cell Dev Biol 20, 781–810 (2004).

50. Masckauchán, T. N. H., Shawber, C. J., Funahashi, Y., Li, C.-M. & Kitajewski, J. Wnt/beta-catenin signaling induces proliferation, survival and interleukin-8 in human endothelial cells. Angiogenesis 8, 43–51 (2005).

51. Sutherland, C. What are the bona fide GSK3 substrates? Int J Alzheimers Dis 2011, (2011).

52. Nie, X., Liu, H., Liu, L., Wang, Y. D. & Chen, W. D. Emerging Roles of Wnt Ligands in Human Colorectal Cancer. Front Oncol 10, 1–13 (2020).

53. Wang, R. & Tian, B. APAlyzer: A bioinformatics package for analysis of alternative polyadenylation isoforms. Bioinformatics 36, 3907–3909 (2020).

54. Ge, S. et al. Interleukin 17 receptor A modulates monocyte subsets and macrophage generation in vivo. PLoS One 9, (2014).

55. Yan, C. et al. IL-17R deletion predicts high-grade colorectal cancer and poor clinical outcomes. Int J Cancer 145, 548–558 (2019).

56. Zykova, T. et al. Targeting prpk function blocks colon cancer metastasis. Mol Cancer Ther 17, 1101–1113 (2018).

57. Gross, J. C., Chaudhary, V., Bartscherer, K. & Boutros, M. Active Wnt proteins are secreted on exosomes. Nat Cell Biol 14, 1036–1045 (2012).

58. Shi, M. et al. ALYREF mainly binds to the 5′ and the 3′ regions of the mRNA in vivo. Nucleic Acids Res 45, 9640–9653 (2017).

59. Zhou, H. et al. IRAK2 directs stimulus-dependent nuclear export of inflammatory mRNAs. Elife 6, 1–25 (2017).

60. Xu, G. et al. Transcriptome and targetome analysis in MIR155 expressing cells using RNA-seq. Rna 16, 1610–1622 (2010).

61. Pyo, J.-S. S., Park, M. J. & Kim, C.-N. N. TPL2 expression is correlated with distant metastasis and poor prognosis in colorectal cancer. Hum Pathol 79, 50–56 (2018).

62. Chan, S. L. et al. CPSF30 and Wdr33 directly bind to AAUAAA in mammalian mRNA 3′ processing. Genes Dev 28, 2370–2380 (2014).

63. Makler, A. & Narayanan, R. Mining exosomal genes for pancreatic cancer targets. Cancer Genomics Proteomics 14, 161–172 (2017).

64. Ciavarella, S. et al. A peculiar molecular profile of umbilical cord-mesenchymal stromal cells drives their inhibitory effects on multiple myeloma cell growth and tumor progression. Stem Cells Dev 24, 1457–1470 (2015).

65. Zhao, S. et al. Exploration of a Novel Prognostic Risk Signature and Its Effect on the Immune Response in Nasopharyngeal Carcinoma. Front Oncol 11, 1–12 (2021).

66. Zhang, Y., Feng, J., Cui, J., Yang, G. & Zhu, X. Pre-B cell leukemia transcription factor 3 induces inflammatory responses in human umbilical vein endothelial cells and murine sepsis via acting a competing endogenous RNA for high mobility group box 1 protein. Mol Med Rep 17, 5805–5813 (2018).

67. Liu, Y., Ao, X., Zhou, X., Du, C. & Kuang, S. The regulation of PBXs and their emerging role in cancer. J Cell Mol Med 26, 1363–1379 (2022).

68. Han, H. B. et al. PBX3 promotes migration and invasion of colorectal cancer cells via activation of MAPK/ERK signaling pathway. World J Gastroenterol 20, 18260–18270 (2014).

69. Kühnemuth, B. & Michl, P. The role of CUX1 in antagonizing NF-κB signaling in TAMs. Oncoimmunology 3, (2014).

70. Ng, L. F. et al. WNT signaling in disease. Cells 8, (2019).

71. Wang, L. et al. Oxidized phospholipids are ligands for LRP6. Bone Res 6, (2018).

72. Hamdollah Zadeh, M. A., et al. Alternative splicing of TIA-1 in human colon cancer regulates VEGF isoform expression, angiogenesis, tumour growth and bevacizumab resistance. Mol Oncol 9, 167–178 (2015).

73. Saito, K., Chen, S., Piecyk, M. & Anderson, P. TIA-1 regulates the production of tumor necrosis factor alpha in macrophages, but not in lymphocytes. Arthritis Rheum 44, 2879–2887 (2001).

74. Serra, M., Columbano, A., Ammarah, U., Mazzone, M. & Menga, A. Understanding Metal Dynamics Between Cancer Cells and Macrophages: Competition or Synergism? Front Oncol 10, 1–16 (2020).

75. Micaroni, M. et al. Rab6a/a’ Are Important Golgi Regulators of Pro-Inflammatory TNF Secretion in Macrophages. PLoS One 8, (2013).

76. Zhou, X. et al. Na+/H+-exchanger family as novel prognostic biomarkers in colorectal cancer. J Oncol 2021, (2021).

77. Cowper, A. E. et al. Serine-Arginine (SR) Protein-like Factors That Antagonize Authentic SR Proteins and Regulate Alternative Splicing. Journal of Biological Chemistry 276, 48908–48914 (2001).

78. Ho, T. H. et al. Muscleblind proteins regulate alternative splicing. EMBO Journal 23, 3103– 3112 (2004).

79. Batra, R. et al. Loss of MBNL Leads to Disruption of Developmentally Regulated Alternative Polyadenylation in RNA-Mediated Disease. Mol Cell 157, 1644–1656 (2015).

80. Allen, W. E., Zicha, D., Ridley, A. J. & Jones, G. E. A role for Cdc42 in macrophage chemotaxis. Journal of Cell Biology 141, 1147–1157 (1998).

81. Lee, D. J., Cox, D., Li, J. & Greenberg, S. Rac1 and Cdc42 are required for phagocytosis, but not NF-κB-dependent gene expression, in macrophages challenged with Pseudomonas aeruginosa. Journal of Biological Chemistry 275, 141–146 (2000).

82. Sica, A., Schioppa, T., Mantovani, A. & Allavena, P. Tumour-associated macrophages are a distinct M2 polarised population promoting tumour progression: Potential targets of anti-cancer therapy. Eur J Cancer 42, 717–727 (2006).

83. Bao, X. et al. Integrated analysis of single-cell RNA-seq and bulk RNA-seq unravels tumour heterogeneity plus M2-like tumour-associated macrophage infiltration and aggressiveness in TNBC. Cancer Immunol Immunother 70, 189–202 (2021).

84. Mombelli, S. et al. IL-17A and its homologs IL-25/IL-17E recruit the c-RAF/S6 kinase pathway and the generation of pro-oncogenic LMW-E in breast cancer cells. Sci Rep 5, 1–10 (2015).

85. Facchin, S. et al. Functional homology between yeast piD261/Bud32 and human PRPK: Both phosphorylate p53 and PRPK partially complements piD261/Bud32 deficiency. FEBS Lett 549, 63–66 (2003).

86. Morris, A. R. et al. Alternative cleavage and polyadenylation during colorectal cancer development. Clinical Cancer Research 18, 5256–5266 (2012).

87. Takagaki, Y., Seipelt, R. L., Peterson, M. L. & Manley, J. L. The polyadenylation factor CstF-64 regulates alternative processing of IgM heavy chain pre-mRNA during B cell differentiation. Cell 87, 941–952 (1996).

88. Amara, S. G., Evans, R. M. & Rosenfeld, M. G. Calcitonin/calcitonin gene-related peptide transcription unit: tissue-specific expression involves selective use of alternative polyadenylation sites. Mol Cell Biol 4, 2151–2160 (1984).

89. Edwalds-Gilbert, G., Veraldi, K. L. & Milcarek, C. Alternative poly(A) site selection in complex transcription units: Means to an end? Nucleic Acids Res 25, 2547–2561 (1997).

90. Berg, M. G. et al. U1 snRNP determines mRNA length and regulates isoform expression. Cell 150, 53–64 (2012).

91. Lou, H., Neugebauer, K. M., Gagel, R. F. & Berget, S. M. Regulation of Alternative Polyadenylation by U1 snRNPs and SRp20. Mol Cell Biol 18, 4977–4985 (1998).

92. So, B. R. et al. A Complex of U1 snRNP with Cleavage and Polyadenylation Factors Controls Telescripting, Regulating mRNA Transcription in Human Cells. Mol Cell 76, 590–599.e4 (2019).

93. Howard, J. M. & Sanford, J. R. The RNAissance family: SR proteins as multifaceted regulators of gene expression. Wiley Interdiscip Rev RNA 6, 93–110 (2015).

94. Martin, M. Cutadapt removes adapter sequences from high-throughput sequencing reads. EMBnet 7, 2803–2809 (2011).

95. Langmead, B., Trapnell, C., Pop, M. & Salzberg, S. L. Ultrafast and memory-efficient alignment of short DNA sequences to the human genome. Genome Biol 10, (2009).

96. Trapnell, C., Pachter, L. & Salzberg, S. L. TopHat: discovering splice junctions with RNA-Seq. Bioinformatics 25, 1105–1111 (2009).

97. Dobin, A. STAR manual 2.4.0.1. 1–33 (2014) doi:10.1093/bioinformatics/bts635.

98. Dobin, A. et al. Mapping RNA-seq Reads with STAR. Curr Protoc Bioinformatics 51, 586–597 (2016).

99. Li, H. et al. The Sequence Alignment/Map format and SAMtools. Bioinformatics 25, 2078– 2079 (2009).

100. Anders, S., Pyl, P. T. & Huber, W. HTSeq-A Python framework to work with high-throughput sequencing data. Bioinformatics 31, 166–169 (2015).

101. Love, M., Anders, S. & Huber, W. Analyzing RNA-seq data with DESeq2. Bioconductor 2, 1–63 (2017).

102. Wang, R. & Tian, B. APAlyzer: A bioinformatics package for analysis of alternative polyadenylation isoforms. Bioinformatics 36, 3907–3909 (2020).

103. Schindelin, J. et al. Fiji: An open-source platform for biological-image analysis. Nat Methods 9, 676–682 (2012).

104. Livak, K. J. & Schmittgen, T. D. Analysis of relative gene expression data using real-time quantitative PCR and the 2-ΔΔCT method. Methods 25, 402–408 (2001).

105. Fischl, H. et al. hnRNPC regulates cancer-specific alternative cleavage and polyadenylation profiles. Nucleic Acids Res 47, 7580–7591 (2019).

106. Pereira-Castro, I. et al. MCL1 alternative polyadenylation is essential for cell survival and mitochondria morphology. Cellular and Molecular Life Sciences 79, (2022).

107. Nojima, T. et al. Mammalian NET-seq reveals genome-wide nascent transcription coupled to RNA processing. Cell 161, 526–540 (2015).

108. Dobin, A. et al. STAR: Ultrafast universal RNA-seq aligner. Bioinformatics 29, 15–21 (2013).

109. Love, M. I., Huber, W. & Anders, S. Moderated estimation of fold change and dispersion for RNA-seq data with DESeq2. Genome Biol 15, 1–21 (2014).

110. The Cancer Genome Atlas. https://cancergenome.nih.gov/.

